# The dual ubiquitin binding mode of SPRTN secures rapid spatiotemporal proteolysis of DNA-protein crosslinks

**DOI:** 10.1101/2024.11.26.625361

**Authors:** Wei Song, Yichen Zhao, Annamaria Ruggiano, Christina Redfield, Joseph A Newman, Xiaosheng Zhu, Abimael Cruz-Migoni, Rebecca Roddan, Peter McHugh, Paul Elliott, Kristijan Ramadan

## Abstract

DNA-protein crosslinks (DPCs) are endogenous and chemotherapy-induced genotoxic DNA lesions and, if not repaired, lead to embryonic lethality, neurodegeneration, premature ageing, and cancer. DPCs are heavily polyubiquitinated, and the SPRTN protease and 26S proteasome emerged as two central enzymes for DPC proteolysis. The proteasome recognises its substrates by their ubiquitination status. How SPRTN protease, an essential enzyme for DPC proteolysis, achieves specificity for DPCs still needs to be discovered. We found that the N-terminal SPRTN catalytic region (SprT) possesses a ubiquitin-binding domain named the <u>U</u>biquitin interface of <u>S</u>prT <u>D</u>omain (USD). Using multiple biochemical, biophysical, and structural approaches, we reveal that USD binds ubiquitin chains. SPRTN binding to ubiquitin chains via USD leads to ∼ 67-fold higher activation of SPRTN proteolysis towards polyubiquitinated DPCs than the unmodified DPCs. This study reveals the ubiquitination of DPCs is the key signal for SPRTN’s substrate specificity and rapid proteolysis.

## INTRODUCTION

DPCs are bulky cytotoxic DNA lesions induced by the covalent attachment of chromatin-associated proteins to DNA in the presence of crosslinking agents.^1–7^ DPCs form with exposure to endogenous metabolic products, such as aldehydes and chemotherapy drugs, such as Topoisomerase inhibitors and Oxaliplatin. DPCs can also form due to the physiological or abortive actions of DNA metabolic enzymes. Examples include the binding of HMCES (5hmC-binding, embryonic stem-cell-specific protein) to sites of abasic lesions, which protects single-stranded DNA from breaks during replication, and the covalent trapping of topoisomerases at endogenous DNA lesions.^8–10^ DPCs disrupt chromatin structures and suppress the progression of DNA replication and transcription due to their bulkiness. If not repaired, DPCs lead to embryonic lethality, premature ageing, neurodegeneration and cancer.^11–18^ Therefore, understanding the DPC repair mechanisms is essential for preventing these diseases and improving cancer therapies.

The study of DPC repair has been significantly boosted over the last decade by discovering a specialised replication-coupled DPC proteolysis repair pathway in yeast and vertebrates.^11,12,19–23^ During DNA replication, the metalloproteases Wss1 in yeast and its counterpart SPRTN (also known as DVC1) in vertebrates interact with the replisome and travel alongside DNA replication.^19,24–26^ As a protease, Wss1 and SPRTN cleave the protein component of the DPC, leaving a remnant peptide that enables the progression of the DNA replication fork via translesion synthesis polymerases.^20,27^ Notably, SPRTN knock-out causes early embryonic lethality in mice, and its biallelic mutations or down-regulation causes Ruijs-Aalfs Syndrome (RJALS) in humans or RJALS-like phenotype in mice, respectively.^11,12^ RJALS is characterised by chromosomal instability, accelerated ageing/segmental progeria, and early-onset hepatocellular carcinoma.^11,12^ SPRTN is a DNA-dependent metalloprotease but lacks substrate specificity.^19,22,23^ Therefore, SPRTN must be tightly regulated to avoid unintended cleavage of the DNA replication machinery or unspecific proteolysis of functional chromatin proteins.

In recent years, several levels of SPRTN regulation have been discovered.^28,29^ SPRTŃs activity strictly depends on DNA binding. It is achieved by two DNA binding interfaces, the Zn-binding domain (ZBD) and the basic DNA binding region (BR) in the N-terminal part of SPRTN, close to the active centre of its protease activity.^3,30^ SPRTN’s activity is highly unselective *in vitro*, where virtually any DNA-bound protein can be proteolysed, including SPRTN itself.^19,22,23^ This autocleavage activity is deemed to be a mechanism of self-inactivation.^22,31^ Whether SPRTŃs activity is targeted to substrates or autocleavage is determined by the nature of the activating DNA, although controversies exist. Some studies have demonstrated that short single-stranded oligonucleotides (60 mer) and circular single-stranded (ss) DNA plasmids induce efficient SPRTN protease activity towards substrates.^22^ In contrast, double-stranded (ds) DNA mainly governs its activity for auto-cleavage.^22^ Other studies claim that dsDNA also activates SPRTN towards substrates.^19,23^ Regardless, the best activating DNA structure *in vitro* is the ss/dsDNA junction (dsDNA with overhang).^3^ During DNA replication, a similar DNA structure would result from uncoupling between the DPC-stalled DNA polymerase and the CMG helicase, which might bypass the DPC lesion.^32^ The second level of SPRTN regulation relies on ubiquitination, but the mechanism is largely elusive and controversial. SPRTN possesses a UBZ domain that directly binds to ubiquitin. Data from *in vitro* and a cell-free system suggest that SPRTN processing of substrates occurs independent of their ubiquitination.^3,19,22,23,27,30^ A partial explanation is that the SPRTN UBZ domain may not directly bind to DPC substrates; instead, it may engage with other ubiquitinated proteins in the vicinity of DPCs.^27^ One candidate is PCNA, a ubiquitination target in response to DNA damage.^33^ This leads to a widely accepted dogma in the field that SPRTN is involved in processing non-ubiquitinated DPCs and that the 26S proteasome is the central protease for processing ubiquitinated DPCs.^8,28,34^ However, other studies reported that SPRTN localization to formaldehyde-induced nuclear repair foci relies on a viable ubiquitination signal and a functional UBZ domain.^35,36^ What’s more, depletion of SPRTN induced the accumulation of ubiquitinated DPCs in cells,^35^ indicating that, most likely, ubiquitinated DPCs are also substrates for SPRTN proteolysis.

Ubiquitin also seems to regulate SPRTN protease activity. It has been reported that adding ubiquitin to the reaction *in vitro* facilitates SPRTN autocleavage and proteolysis towards Top2α, but there is no further understanding of this effect.^23^ Recent data suggest that monoubiquitination on SPRTN is a regulatory mechanism for SPRTN inactivation and consequent degradation by auto-cleavage and proteasomal degradation.^31^

Altogether, varying evidence suggests that the regulation of SPRTN recruitment and proteolysis of DPCs depends on ubiquitination at/around DNA damage sites. However, the exact role of SPRTN in processing modified DPCs, including ubiquitinated substrates, has yet to be characterised and understood.

Thus, we hypothesised that ubiquitin and the ubiquitin signal have a much more direct role in regulating SPRTN-dependent DPC proteolysis and might contribute to SPRTN’s specificity towards substrates. We first engineered a high-sensitivity fluorescence-based assay to monitor SPRTN proteolysis in combination with ubiquitin *in vitro*. Using biophysical methods to address protein-protein interactions, including ITC and NMR, we discovered that SPRTN has dual ubiquitin-binding properties that govern its rapid proteolysis of polyubiquitinated substrates, including DPCs. Specifically, the C-terminal UBZ domain of SPRTN, which has a high affinity to ubiquitin, acts as a general sensor for ubiquitinated substrates. In contrast, the here identified N-terminal ubiquitin-binding interface (USD), located on the SprT domain, allows SPRTN to selectively bind to polyubiquitinated substrates in an avidity manner and, consequently, facilitates their rapid proteolysis. The polyubiquitin chains on DPCs potentiate the enzymatic activity of SPRTN, making the substrate processing remarkably faster than non-ubiquitinated substrates. Our work has resolved a long-standing question in the field, namely, how SPRTN gains its substrate specificity and further demonstrates that polyubiquitin chains on DPCs are essential catalysts for SPRTN, revealing an exquisitely fine-tuned regulation of SPRTN proteolysis.

## RESULTS

### SPRTN proteolysis towards the substrate is activated by single-stranded (ss) and double-stranded (ds) DNA

To gain mechanistic insights into SPRTN-dependent substrate proteolysis, we chose histone H1 as a model substrate. Histone H1 has been widely used to study SPRTN proteolysis *in vitro*.^3,30^ To confirm that histone H1 is a relevant substrate for SPRTN *in vivo*, we depleted SPRTN in U2OS cells (Figure S1A). SPRTN depletion by siRNA caused increased covalent attachment of histone H1 to genomic DNA when analysed by the RADAR assay, a well-recognised method to measure DNA-protein crosslinks^37,38^ (Figure S1B). We also found that histone H1 is an abundantly crosslinked protein in response to formaldehyde (Figure S1C). Interestingly, formaldehyde treatment further induced the accumulation of high molecular weight species which react with an anti-H1.0 specific antibody (Figure S1C), most likely post-translationally modified histone H1^35,39^.

To monitor SPRTŃs proteolytic activity on H1 and bypass the limitations of western blot detection of its cleavage products^3,22,30^, we developed a fluorescence-based *in vitro* assay (Figure 1A). Cy5-labelled N-or C-terminal histone H1 (Cy5-N-H1 and H1-C-Cy5 hereafter) were generated to monitor SPRTN’s enzymatic activity (Figure 1A, lower panel).

**Figure 1.**
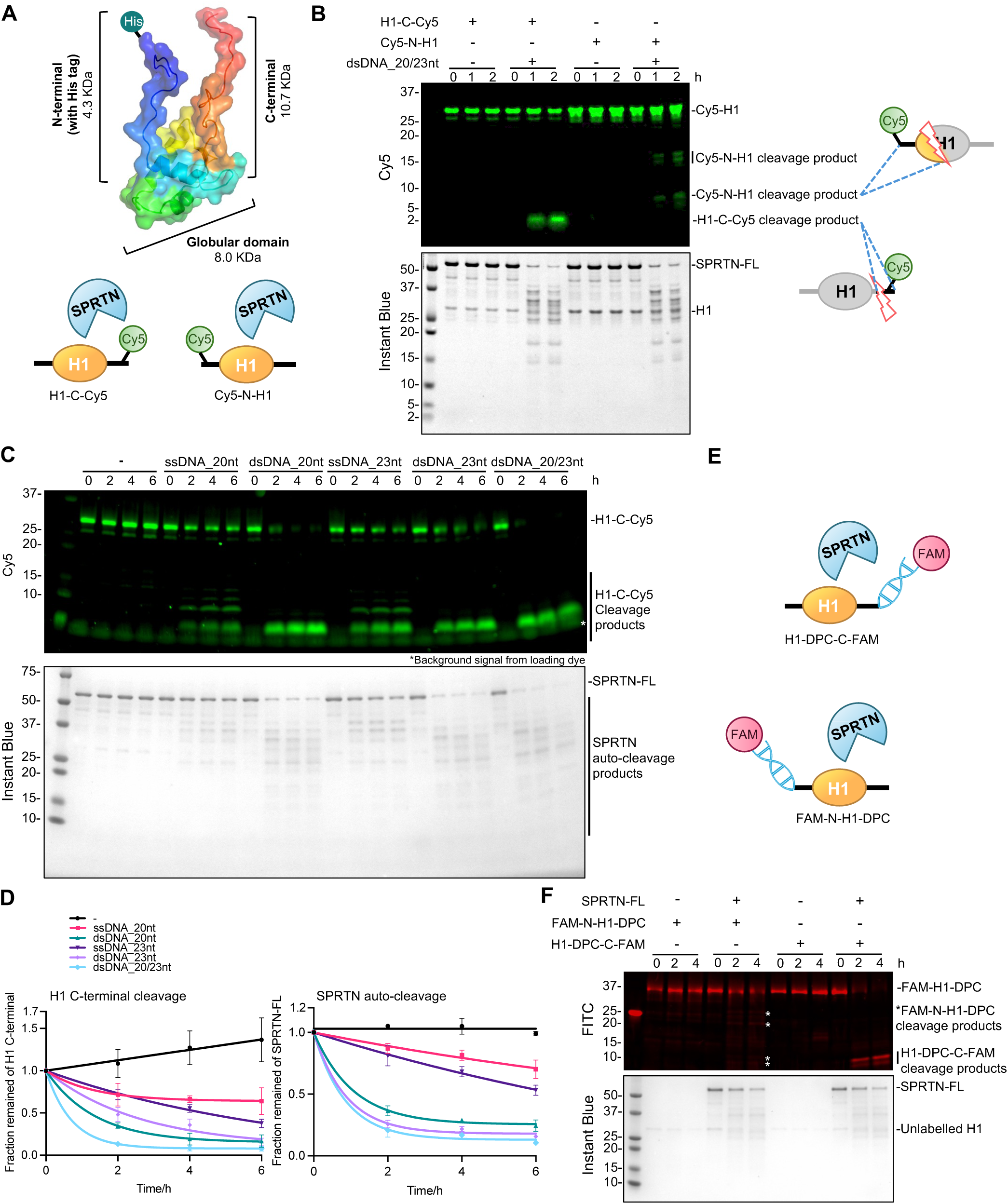
SPRTN proteolysis is activated by both ssDNA and dsDNA. **(A)** Design of the Cy5-labelled H1 substrates (Cy5-C-H1 and Cy5-N-H1) for SPRTN cleavage assay. The size of the H1 N-terminal (including the His tag), C-terminal and globular domain is indicated, respectively. **(B)** SPRTN cleavage assay towards the model H1 substrates. Recombinant full-length SPRTN (2 uM) and Cy5-labelled H1 substrates (H1-C-Cy5 or Cy5-N-H1, 1 uM) were incubated in the absence or presence of dsDNA_20/23nt (2.7 uM) with the indicated time at 30°C. The reaction was analysed by SDS-PAGE followed by Cy5-scanning on Typhoon FLA 9500 (GE Healthcare) and Instant Blue staining. Representative figure from 3 repeats. **(C)** SPRTN kinetics towards H1-C-Cy5. Recombinant full-length SPRTN (2 µM) and H1-C-Cy5 (1 µM) were incubated with indicated ssDNAs or dsDNAs (2.7 µM) at the indicated time at 30°C. The reaction was analysed by SDS-PAGE followed by Cy5-scanning on an iBright 1500 imaging system (Invitrogen) and Instant Blue staining. Representative figure from 3 repeats. **(D)** Kinetics of the full-length H1-C-Cy5 substrate (C-terminal cleavage rate) and the full-length SPRTN (auto-cleavage rate) from Figure 1C. Cy5 and SPRTN-FL signals were analysed by the iBright Analysis Software (Invitrogen). Kinetic data were fitted with one phase exponential decay - least squares fit (Prism). n=3. Error bar, SD. **(E)** Design of the FAM-dsDNA labelled H1-DPC substrates (H1-DPC-C-FAM and FAM-N-H1-DPC) for SPRTN cleavage assay. **(F)** SPRTN cleavage assay towards the model H1-DPC substrates. Recombinant full-length SPRTN (2 µM) and the model H1-DPC substrates (H1-DPC-C-FAM or FAM-N-H1-DPC, ∼0.1 µM) were incubated with the indicated time at 30°C. The reaction was analysed by SDS-PAGE followed by FITC-scanning on an iBright 1500 imaging system (Invitrogen) and Instant Blue staining. Representative figure from 2 repeats. See also Figure S1.

As SPRTŃs activity is strictly DNA-dependent, we first used dsDNA with a 3-nt overhang (dsDNA 20/23nt) as it was reported as the best DNA source to activate SPRTN proteolysis.^3^ It was designed based on known ssDNA sequences that do not precipitate histone H1 when used in SPRTN proteolysis assay.^40^ Importantly, our Cy5-fluorescence-based assays provided a higher resolution of histone H1 cleavage products by SPRTN proteolysis than previously reported^3,22,30^ (Figure 1B). SPRTN was shown to cleave mostly unstructured regions of the core histones, namely the histone tails.^19^ Likewise, SPRTN preferably cleaved the more disordered C-terminal (Figure 1B). The cleavage at the N-terminus of histone H1 was rate-limited. However, multiple cleavage products were observed, which indicate that SPRTN cleaves not only the unstructured N-terminal tail (N-terminal tail is 4.3 kDa including the His tag, Figure 1A) but also structured globular regions of H1 (Figure 1B, the last two lanes), as one of the cleavages products are between 5 and 10 kDa.

There is an ongoing debate regarding how different DNA templates activate SPRTN. A dogma has been accepted that ssDNA activates SPRTN towards the substrate and blunt-ended dsDNA towards auto-cleavage, consequently causing SPRTN inactivation. However, it has been shown that SPRTN cleaves core histones, denatured TOP1/TOP2 and CHK1, in the presence of 100-nt dsDNA with blunt ends.^19,25^ In contrast, other studies demonstrated that SPRTN proteolysis towards histone H1 cannot be activated by blunt-ended or circular dsDNAs (ranging from 60 nt to 5.4 Kb),^3,30^ and that the presence of either ssDNA or dsDNA determines better substrate cleavage or auto-cleavage, respectively.^22^ To clear up this controversy, we took advantage of our assay, where cleaving the C-terminal fluorescently labelled H1 (H1-C-Cy5, Figure 1C) provides a clear and robust readout of SPRTN activity. Different ssDNAs and dsDNAs were screened based on our fluorescence assay, and both cleavage of histone H1 and auto-cleavage of SPRTN were monitored (Figures 1C and 1D). Results demonstrated that blunt-ended dsDNAs (without overhang) could also activate SPRTN proteolysis, as a very small Cy5-visible cleavage product around 2 KDa appeared, which corresponds to the very end of H1 C-terminal (“dsDNA_20nt”, “dsDNA_23nt”, Figures 1C and S1D). In addition, ssDNA facilitated SPRTN attack into the structured region of histone H1 (“ssDNA_20nt”, ssDNA_23nt”, Figures 1C and S1D) as additional cleavage fragments with a size roughly above 5 KDa appeared (Figure S1D). In line with a previous study,^3^ dsDNA, but not ssDNA, induced strong SPRTN auto-cleavage within 2 hours (Figures 1C and 1D). Still, ssDNA governs the specificity towards substrates over self-cleavage.

Notably, the SPRTN proteolysis kinetics tested by various DNA structures (ss, ds, overhangs) showed the most rapid cleavage towards the C-terminal tail of histone H1 was achieved by the ss/dsDNA overhang (“dsDNA_20/23nt”, Figures 1C, 1D and S1D), which is in agreement with previous reports.^3^ However, although the 3-nt overhang dsDNA structure (20/23nt) was the best activator of SPRTN proteolysis towards both substrate-cleavage and auto-cleavage, the cleavage kinetics were still relatively slow (∼2h), indicating SPRTN is not a very efficient enzyme (Figures 1C and 1D).

A model H1-DPC substrate was also developed where the FAM-labelled dsDNA was covalently attached to either the N- or C-terminal of H1 (Figure 1E). Again, SPRTN proteolysis towards the model H1-DPCs indicated a cleavage pattern similar to free H1 (please compare Figure 1F with Figure 1B).

Altogether, we established a sensitive fluorescence-based assay to monitor SPRTN proteolysis on a genuine DPC substrate, histone H1, *in vitro*. We demonstrated that single-stranded and double-stranded DNA structures activate SPRTN proteolytic activity towards itself and the substrate. In addition, we confirmed that the 3-nt overhang dsDNA structure (20/23nt) was the best activator of SPRTN proteolysis. SPRTN cleaves its substrate and itself on this DNA structure with similar efficiency (Figures 1C and 1D, last four lanes), suggesting its inactivation during substrate processing and no substrate specificity.

### Ubiquitin chains rapidly activate SPRTN proteolysis

The slow kinetics of substrate cleavage (Figures 1 and S1) suggest that SPRTN is a very inefficient enzyme. SPRTN is also a promiscuous enzyme that cleaves many DNA-bound proteins.^3,19,22,25,30^ This is counterintuitive, considering its role as a DNA replication coupled enzyme for the timely resolution of DPCs in front of the DNA replication fork.^19,21,25^ The replication fork is enriched with the replisome, whose unspecific cleavage would harm the proliferative cell.^41^ All these facts suggest that additional factors must regulate SPRTN enzymatic activity and specificity. As SPRTN possesses the Ub-binding domain Zn finger (UBZ) at its C-terminal (Figure 2A) and DPCs are heavily ubiquitinated,^27,35^ we hypothesised that ubiquitin or ubiquitination on its substrates could play a regulatory role for SPRTN proteolysis and specificity.

**Figure 2.**
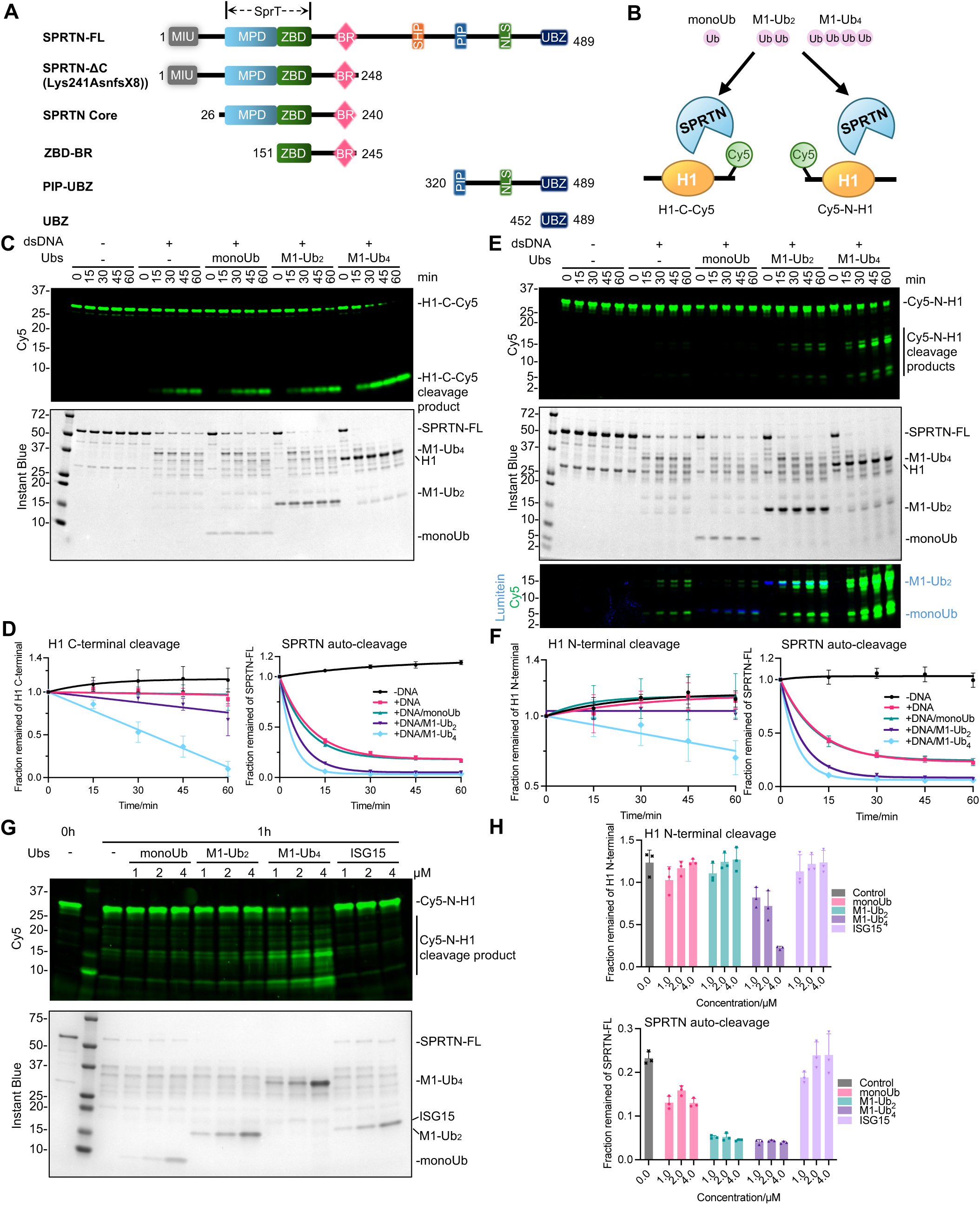
Ubiquitin chains rapidly activate SPRTN proteolysis. **(A)** Schematics of SPRTN domain structure. MIU, Motif Interacting with Ub-binding domain; SprT, the metalloprotease domain similar to that of the *E. coli* SprT protein; BR, basic region; PIP, PCNA interaction peptide; SHP, p97 or VCP-binding motif; SIM, SUMO interaction motif; VIM; VCP interaction motif; UBZ, ubiquitin-binding zinc finger. **(B)** Schematics of Ub titration to SPRTN cleavage assay with Cy5-labelled H1 substrates (Cy5-C-H1 or Cy5-N-H1). **(C)** SPRTN cleavage assay towards H1-C-Cy5 in the presence of Ubs. Recombinant full-length SPRTN (2 µM) was incubated with H1-C-Cy5 (1 µM) in the absence or presence of dsDNA_20/23nt (2.7 µM) in combination with Ubs (monoUb, M1-Ub_2_, M1-Ub_4_, all at 2 µM) with the indicated time at 30°C. Representative figure from 3 repeats. **(D)** Cleavage kinetics of the full-length H1-C-Cy5 substrate (C-terminal cleavage rate) and the full-length SPRTN (auto-cleavage rate) from Figure 2C. n=3. Error bar, SD. **(E)** SPRTN cleavage assay towards Cy5-N-H1 in the presence of Ubs. Recombinant full-length SPRTN (2 µM) was incubated with Cy5-N-H1 (1 µM) in the absence or presence of dsDNA_20/23nt (2.7 µM) in combination with Ubs (monoUb, M1-Ub_2_, M1-Ub_4_, all at 2 µM) with the indicated time at 30°C. The overlay of Cy5 and Lumitein signals (both with stronger contrast) is also displayed to show the similar size of the H1 cleavage product and monoUb. Representative figure from 4 repeats. **(F)** Cleavage kinetics of the full-length Cy5-N-H1 substrate (N-terminal cleavage rate) and the full-length SPRTN (auto-cleavage rate) from Figure 2E. n=4. Error bar, SD. **(G)** SPRTN cleavage assay towards Cy5-N-H1 with Ubs or ISG15. Recombinant full-length SPRTN (2 µM) and Cy5-N-H1 (1 µM) were incubated with indicated Ubs (monoUb, M1-Ub_2_, M1-Ub_4_) or ISG15 at various concentrations (1-4 µM) in the presence of dsDNA_20/23nt (2.7 µM) for 1h at 30°C. The reactions were analysed with SDS-PAGE followed by Cy5-scanning on an iBright 1500 imaging system (Invitrogen) and Instant Blue staining. Representative figure from 3 repeats. **(H)** Quantification of the signal from the full-length Cy5-N-H1 substrate (N-terminal cleavage rate) and the full-length SPRTN (auto-cleavage rate) from Figure 2G. n=3. Error bar, SD. The reactions from Figures 2C and 2E were analysed with SDS-PAGE followed by Cy5-scanning on Typhoon FLA 9500 (GE Healthcare) and Instant Blue staining. Cy5 signals from Figures 2C and 2E were analysed by ImageJ. Cy5 signals from Figure 2G were analysed by the iBright Analysis Software (Invitrogen). SPRTN-FL signals from Figures 2C, 2E and 2G were analysed by the iBright Analysis Software (Invitrogen). Kinetic data from Figure 2D and 2F were fitted with one phase exponential decay - least squares fit (Prism), respectively. See also Figure S2.

Using the above-established H1-C-Cy5 as a substrate for SPRTN proteolysis in the presence of 3-nt overhang dsDNA (20/23nt), we added either monoUb, M1-linked diUb (M1-Ub_2_) or M1-linked tetraUb (M1-Ub_4_) to test our hypothesis (Figure 2B). Interestingly, the addition of M1-Ub_2_, but not monoUb, in the reaction weakly increased the cleavage capacity of SPRTN on histone H1 (Figures 2C and 2D), whereas the addition of M1-Ub_4_ in the reaction induced strong activation of SPRTN proteolytic activity towards itself and the substrate histone H1. Almost half of the C-terminal tail from H1 was cleaved in 30 min and 100% in 60 min (Figures 2C and 2D). As ubiquitin itself could not activate SPRTN without DNA in the reaction (Figure S2A), this indicates that ubiquitin activation is the second tier in regulating SPRTN proteolysis. At the same time, DNA is still a prerequisite for SPRTN activation.

A similar Ub-chain activation effect was observed when Cy5-N-H1 was used as the substrate (Figures 2E and 2F). M1-Ub_4_ significantly enhanced SPRTN proteolysis into the structured histone H1 globular domain, as visible by the decreased full-length H1 and the increased levels of multiple lower molecular weight cleavage products (Figure 2E, the last four lanes). Ubiquitin chain length was essential to sufficiently activate SPRTN, as monoUb or M1-Ub_2_ at higher concentrations could not efficiently activate SPRTN towards Cy5-N-H1 compared to M1-Ub_4_ at a lower concentration (Figures 2G and 2H).

Notably, the ubiquitin-activation of SPRTN protease was not linkage-specific as all eight diUb chains, including the M1, K6, K11, K27, K29, K33, K48, and K63 linkages, could enhance SPRTN activation, among which M1-Ub_2_ displays the best activation on SPRTN proteolysis (Figures S2B and S2C). Furthermore, all tested tetraUb chains promoted C-terminal Histone H1 cleavage in the first 15 min, but K6- and K63-Ub_4_ cleave the C-terminal tail entirely in the first 15 min when compared to other tetraUb chains (Figure S2D).

It’s also worth noting that this activation effect was Ub-specific, as the addition of the Ub-like protein ISG15 in the reaction, which resembles the structure of the linear diUb molecule, did not activate SPRTN (Figures 2G, 2H and S2B, S2C, S2E, S2F). The addition of other Ub-like proteins, such as SUMO1, SUMO2, SUMO3, SUMO2 chains (2-7), or NEDD8 to the reaction did not stimulate SPRTN proteolytic activity (Figures S2E and S2F). Altogether, these results showed that Ub chains, regardless of their linkage-type, activate SPRTN protease activity. This effect is more pronounced if the Ub chains are longer (see Figures 4E and 4F).

### The N-terminal core region of SPRTN is sufficient for its ubiquitin-dependent activation

SPRTN contains a C-terminal UBZ domain (Figure 2A), essential for SPRTN recruitment to DPC substrates and colocalisation at the stalled DNA replication fork.^35,42–44^ To investigate if the UBZ domain is involved in SPRTN activation, we purified a disease-associated variant of SPRTN (SPRTN-ΔC; 1-248 aa) where the C-terminal, including the UBZ domain, was deleted (Figure 2A). Surprisingly, SPRTN-ΔC can still be activated by ubiquitin chains (Figures 3A and 3B) in a dose-dependent manner (Figures 3C and 3D). This set of experiments demonstrates that the UBZ domain is not required for the Ub-mediated activation of SPRTN proteolytic activity.

**Figure 3.**
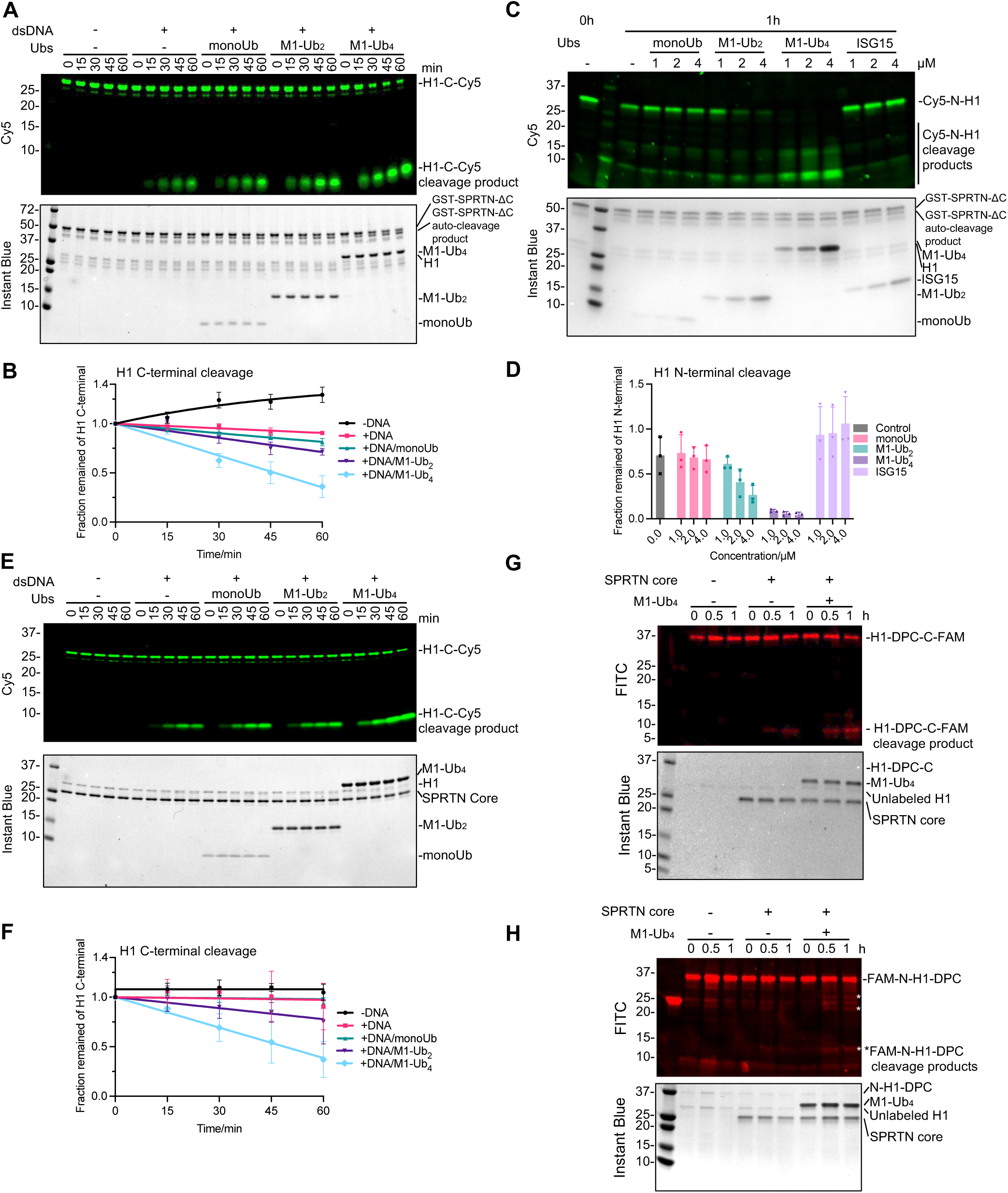
SPRTN core region of SPRTN is sufficient for the rapid activation of SPRTN protease. **(A)** SPRTN-ΔC cleavage assay towards H1-C-Cy5 in the presence of Ubs. Recombinant GST-SPRTN-ΔC (2 µM) and H1-C-Cy5 (1 µM) were incubated in the absence or presence of dsDNA_20/23nt (2.7 µM) in combination with Ubs (monoUb, M1-Ub_2_, M1-Ub_4_, all at 2 µM) with the indicated time at 30°C. Representative figure from 3 repeats. **(B)** Cleavage kinetics of the full-length H1-C-Cy5 substrate (C-terminal cleavage rate) from Figure 3A. n=3. Error bar, SD. **(C)** SPRTN-ΔC cleavage assay towards Cy5-N-Cys in the presence of Ubs or ISG15. Recombinant GST-SPRTN-ΔC (2 µM) and Cy5-N-Cys (1 µM) were incubated with indicated Ubs (monoUb, M1-Ub_2_, M1-Ub_4_) or ISG15 at various concentration (1-4 µM) in the presence of dsDNA_20/23nt (2.7 µM) for 1h at 30°C. Representative figure from 3 repeats. **(D)** Quantification of the signal from the full-length Cy5-N-H1 substrate (N-terminal cleavage rate) from Figure 3C. n=3. Error bar, SD. **(E)** SPRTN core cleavage assay towards H1-C-Cy5 in the presence of Ubs. Recombinant SPRTN core (2 µM) and H1-C-Cy5 (1 µM) were incubated in the absence or presence of dsDNA_20/23nt (2.7 µM) in combination with Ubs (monoUb, M1-Ub_2_, M1-Ub_4_, all at 2 µM) with the indicated time at 30°C. Representative figure from 3 repeats. **(F)** Cleavage kinetics of the full-length H1-C-Cy5 substrate (C-terminal cleavage rate) from Figure 3E. n=3. Error bar, SD. **(G-H)** SPRTN core cleavage assay towards the model H1-DPC substrates with M1-Ub_4_. Recombinant SPRTN core (2 µM) and the model H1-DPC substrates H1-DPC-C-FAM **(G)** or FAM-N-H1-DPC **(H)** (∼0.1 µM) were incubated in the absence or presence of M1-Ub_4_ (2 µM) with the indicated time at 30°C. The reaction was analysed by SDS-PAGE followed by FITC-scanning on an iBright 1500 imaging system (Invitrogen) and Instant Blue staining. Representative figure from 2 repeats. The reactions from Figures 3A and 3E were analysed by SDS-PAGE, followed by Cy5 Scanning on Typhoon FLA 9500 (GE Healthcare) and Instant Blue staining. Cy5 signals were analysed by ImageJ. The reactions from Figure 3C were analysed by SDS-PAGE followed by Cy5-scanning on an iBright 1500 imaging system (Invitrogen) and Instant Blue staining. Cy5 signals were analysed by the iBright Analysis Software (Invitrogen). Kinetic data from Figures 3B and 3F were fitted with a one-phase exponential decay - least squares fit (Prism), respectively. See also Figure S3.

Bioinformatic analysis of the SPRTN protein predicted an additional Motif Interacting with Ubiquitin (MIU) domain located at the first helix of SPRTN (1-20 aa)^11^ (Figure 2A). However, the ubiquitin-binding ability of this putative MIU domain was never experimentally proven. To test the role of the MIU domain in the Ub-dependent activation, we purified a SPRTN variant lacking both the MIU and UBZ domains (Λ1MIU, Λ1C: 26-240 aa, named SPRTN core region hereafter) (Figure 2A). Unexpectedly, the SPRTN core region was still activated by M1-Ub_4_ with kinetics that mimicked SPRTN-ΔC (roughly 60% cleavage within 60 min) (Figures 3E and 3F). The titration of the SPRTN core region, Histone H1 and M1-Ub_4_, in the molar ratio of 2:1:2 caused the most robust activation of the protease, detectable already after only 15 min (Figures S3A and S3B). We also confirmed the ubiquitin-dependent activity of the SPRTN core region towards the model DPC substrates where H1 was covalently attached to FAM-labelled dsDNA in the presence of M1-Ub_4_ (Figures 3G and 3H).

In line with the MIU domain being dispensable in the *in vitro* substrate cleavage assays, isothermal titration calorimetry (ITC) demonstrated that the putative MIU alone could not interact with monoUb or Ub chains (Figure S3C). Competition between the MIU peptide (or GST-MIU) and M1-Ub_4_ in the SPRTN auto-cleavage assay was also not observed (Figures S3D and S3E), indicating that SPRTN MIU does not bind Ub and is not responsible for Ub-dependent SPRTN activation.

Altogether, this suggests that the SPRTN core region alone, which possesses an intrinsic protease domain (MPD), is directly activated by Ub chains, and neither the UBZ nor MIU domain is involved in Ub-dependent SPRTN protease activation.

### The N-terminal SPRTN core region interacts with Ub over a classical Ub Ile44 hydrophobic patch

As the N-terminally located MIU domain neither binds Ub nor is involved in SPRTN activation, we focused on the SPRTN core region (26-240 aa) (Figure 2A). The SPRTN core region is composed of the metalloprotease domain (MPD), where the catalytic active centre is located, the Zn-binding domain (ZBD), and the basic DNA binding region (BR) domain, which is essential for SPRTN protease activation by binding to DNA^3,30^ (Figure 2A). To find out how ubiquitin chains activate SPRTN protease activity, we measured affinity, stoichiometry, and enthalpy changes by ITC between the SPRTN core region and monoUb, M1-Ub_2_, M1-Ub_4_ or M1-Ub_6_ chains (Figures 4A, S4A and Table S1). Enzymatically inactive SPRTN core region (E112Q) for ITC was used to avoid any possible auto-cleavage in the reaction. While binding between the SPRTN core region and monoUb was undetectable by ITC, however, the affinity of the SPRTN core region for M1-Ub_2_ was observed at ∼132 µM, and it further increased to ∼70 µM for M1-Ub_4_ and ∼35 µM for M1-Ub_6_ (Figure 4A and S4A, Table S1). This data strongly suggests that longer Ub chains increase its affinity to the SPRTN core through avidity effects.

**Figure 4.**
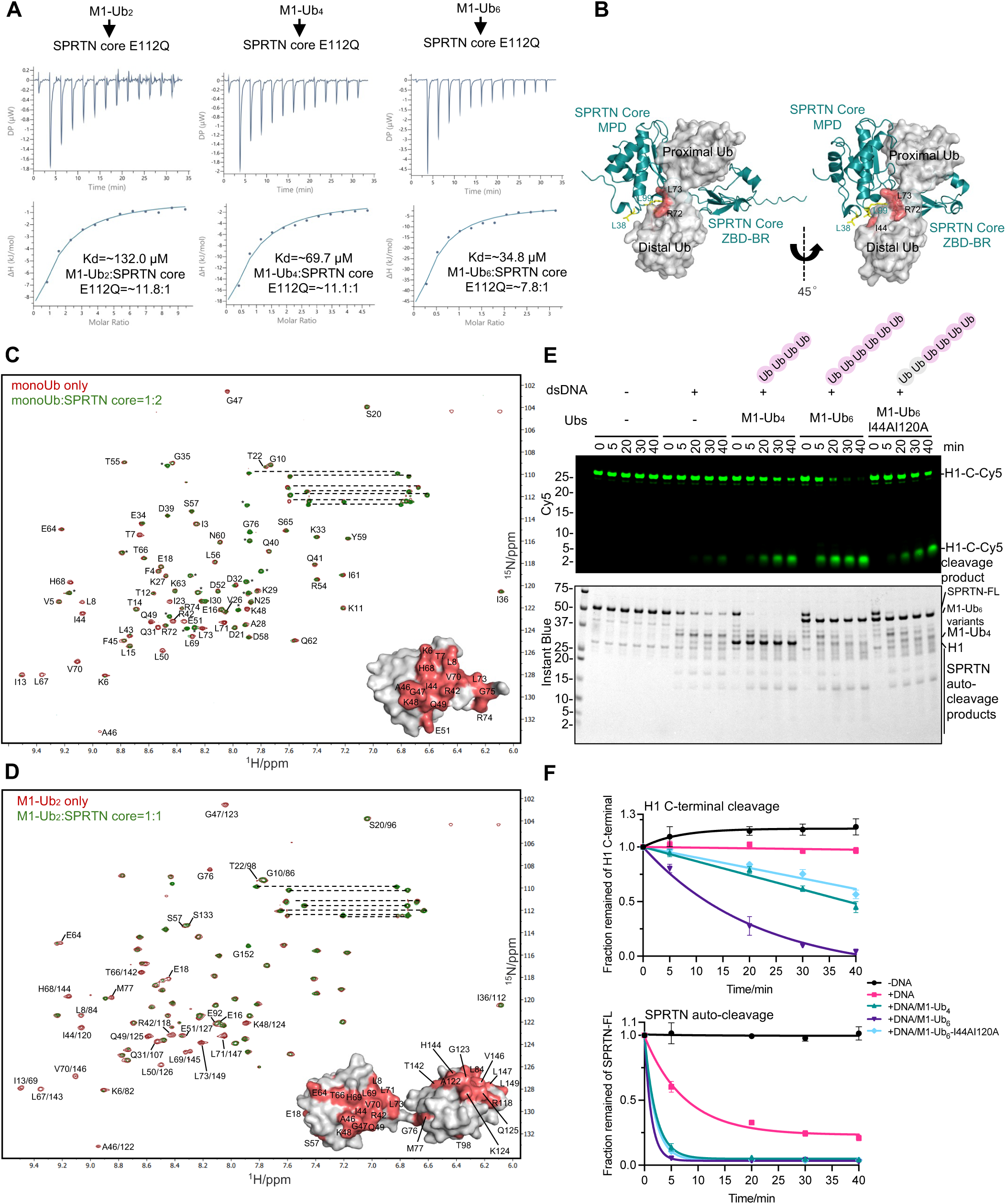
SPRTN core interacts with I44 patch of Ub in an avidity manner. **(A)** Isothermal titration calorimetry (ITC) analysis of the binding between SPRTN core (E112Q) and Ubs (M1-Ub_2_, M1-Ub_4_, M1-Ub_6_). The dissociation constant (Kd) and stoichiometry of binding (N) are indicated here and summarized in Table S1. **(B)** ColaFold prediction of the interaction between SPRTN core and M1-Ub_2_. **(C)** Superimposition of the 950 MHz ^1^H-^15^N HSQC spectra of monoUb alone (in red contours) or with the addition of SPRTN core at a molar ratio of 1:2 (in green contours). Spectra were plotted using the MestReNova software. Residues with complete broadening are highlighted in salmon on the monoUb surface (PDB: 1UBQ). The signals were assigned according to previously reported chemical shifts of ubiquitin (BMRB entry: 17769). *Signals from the N-terminal His tag on monoUb. **(D)** Superimposition of the 950 MHz ^1^H-^15^N HSQC spectra of M1-Ub_2_ alone (in red contours) or with the addition of SPRTN core at a molar ratio of 1:1 (in green contours). Spectra were plotted by using MestReNova software. Residues with complete broadening are highlighted in salmon on the M1-Ub_2_ surface (PDB: 2W9N). The signals were assigned according to previously reported chemical shifts of M1-Ub_2_ (BMRB entry: 26709). **(E)** Validation of M1-Ub_6_ I44 mutant on the effect of SPRTN activation. Recombinant full-length SPRTN (2 µM) and H1-C-Cy5 (1 µM) were incubated in the absence or presence of dsDNA_20/23nt (2.7 µM) in combination with Ubs (M1-Ub_4_, M1-Ub_6_, M1-Ub_6_-I44AI120A, all at 2 µM) with the indicated time at 30°C. The reactions were analysed with SDS-PAGE followed by Cy5-scanning on Typhoon FLA 9500 (GE Healthcare) and Instant Blue staining. Representative figure from 3 repeats. **(F)** Kinetics of the full-length H1-C-Cy5 substrate (C-terminal cleavage rate) and the full-length SPRTN (auto-cleavage rate) from Figure 4E. Cy5 signals were analysed by ImageJ. SPRTN-FL signals were analysed by the iBright Analysis Software (Invitrogen). Kinetic data were fitted with one phase exponential decay - least squares fit (Prism). N=3. Error bar, SD. See also Figure S4.

To narrow down which part of the SPRTN core region binds Ub chains, we purified a SPRTN truncated variant containing only the ZBD-BR domains (Figure 2A). ITC indicated that the ZBD-BR domain did not interact with monoUb, M1-Ub_2_ or M1-Ub_4_ chains (Figure S4A), suggesting that the MPD domain is essential for the Ub-chain binding and SPRTN protease activation. We then used AlphaFold2 to predict the interaction between the SPRTN core region and M1-Ub_2_ (Figures 4B and S4B). AlphaFold2 showed that the SPRTN core region binds M1-Ub_2_ with high confidence (Figure S4B) and predicted a binding interface at the MPD domain that engages with the ubiquitin isoleucine 44 (I44) hydrophobic patch, a crucial surface for the interaction with Ub-binding domains (Figure 4B).

The canonical binding mode of ubiquitin via the I44 patch from the above prediction suggests that there might be a very weak binding between monoUb and the SPRTN core region. Regardless of the undetectable affinity between the SPRTN core and monoUb by ITC (Figure S4A), the analysis by microscale thermophoresis (MST), a more sensitive biophysical method for assessing protein-protein interactions, suggested that the interaction between SPRTN core and monoUb indeed exists, but with an extremely weak affinity at ≥ 769 µM (Figure S4C). We switched to NMR spectroscopy to further explore the physical interaction between the SPRTN core and monoUb, which can extract more detailed information on the binding interfaces, intermolecular affinity, and binding-induced conformational changes for complexes formed in solution. Encouragingly, NMR spectroscopy detected this weak protein-protein interaction in a higher resolution at the single-residue level. The overlaid spectra for the monoUb bound/unbound with SPRTN core confirmed a conserved perturbed surface centred on the I44 patch from ubiquitin (Figure 4C). A similar perturbed pattern from M1-Ub_2_ upon the SPRTN core titration was also illustrated, suggesting the same binding mode between Ub and the SPRTN core region (Figure 4D).

To validate the relevance of the identified interaction between Ub and SPRTN, we mutated I44 to alanine in either the distal Ub (I44A), the proximal Ub (at a position of aa 120; I120) or on both Ub moieties (I44A; I120A) of M1-Ub_2_ and monitored how these mutations affect SPRTŃs affinity to ubiquitin and its proteolytic activity. Specifically, ITC showed that the interaction between the SPRTN core region and the I44A M1-Ub_2_ variants was undetectable, indicating that the affinity is reduced to at least the level of monoUb (Figure S4D). From the Ub-activation assay for SPRTN, it was clear that a single substitution in M1-Ub_2_ was defective in stimulating SPRTN auto-cleavage, while the double mutant M1-Ub_2_-I44A/I120A lost most of its ability to stimulate both auto-cleavage and substrate-cleavage activity of SPRTN (Figures S4E and S4F). We also generated a M1-Ub_6_ variant with mutations in the first two Ub moieties (Figures 4E and 4F). As expected, the cleavage kinetics were faster in the presence of wild-type M1-Ub_6_ than in M1-Ub_4_. More pronouncedly, the M1-Ub_6_-I44AI120A mutant was weaker than the M1-Ub_4_ in the activation on SPRTN (Figures 4E and 4F), further validating the essential role of the Ub I44 patch in the activation effect.

Altogether, these data suggest that the SPRTN core region possesses the intrinsic ability to bind Ub and that Ub binds to the SPRTN core through the classical I44 hydrophobic patch. Ub binding to the SPRTN core is essential for stimulating SPRTN’s protease activity. Most importantly, the affinity of the SPRTN core for Ub is increased with the length of Ub chains, suggesting that SPRTN binds Ub chains with an avidity effect, consequently leading to SPRTN protease activation.

### The SPRTN core region contains an intrinsic Ubiquitin-binding interface

To identify the key residues within the SPRTN core region that engage with the I44 patch of Ub and to have experimental proof that the SPRTN core possesses an intrinsic Ub-binding interface, we carefully looked at the binding interface from the AlphaFold model. We identified L38 and L99 as likely counterparts in the hydrophobic interaction with the I44 patch on Ubiquitin (Figure 4B). SPRTN core variants bearing the L33A or L99A mutation were generated to validate this prediction. We probed their protease activity by comparing it to the SPRTN core wild-type with or without the M1-Ub_4_ chain in the reaction (Figures 5A and 5B; compared the cleavage product ratio indicated below the gel in Figures 5A). The SPRTN core wild-type protease activity was stimulated by M1-Ub_4_ 5-fold when analysed at 60 min. The L38A and L99A variants of the SPRTN core region only possessed residual protease activity compared to the WT. However, the L99A mutant largely lost the ability to be stimulated by M1-Ub_4_ (only 1.5 fold at 60 min), as demonstrated by the reduced ratio between the cleavage products in the presence/absence of M1-Ub_4_ at individual time points, whereas L38A had a minor effect (2.5 fold at 60 min) on Ub-dependent SPRTN activation (Figures 5A and 5B). These data demonstrate the important role of the two residues in the ubiquitin-mediated activation effect.

**Figure 5.**
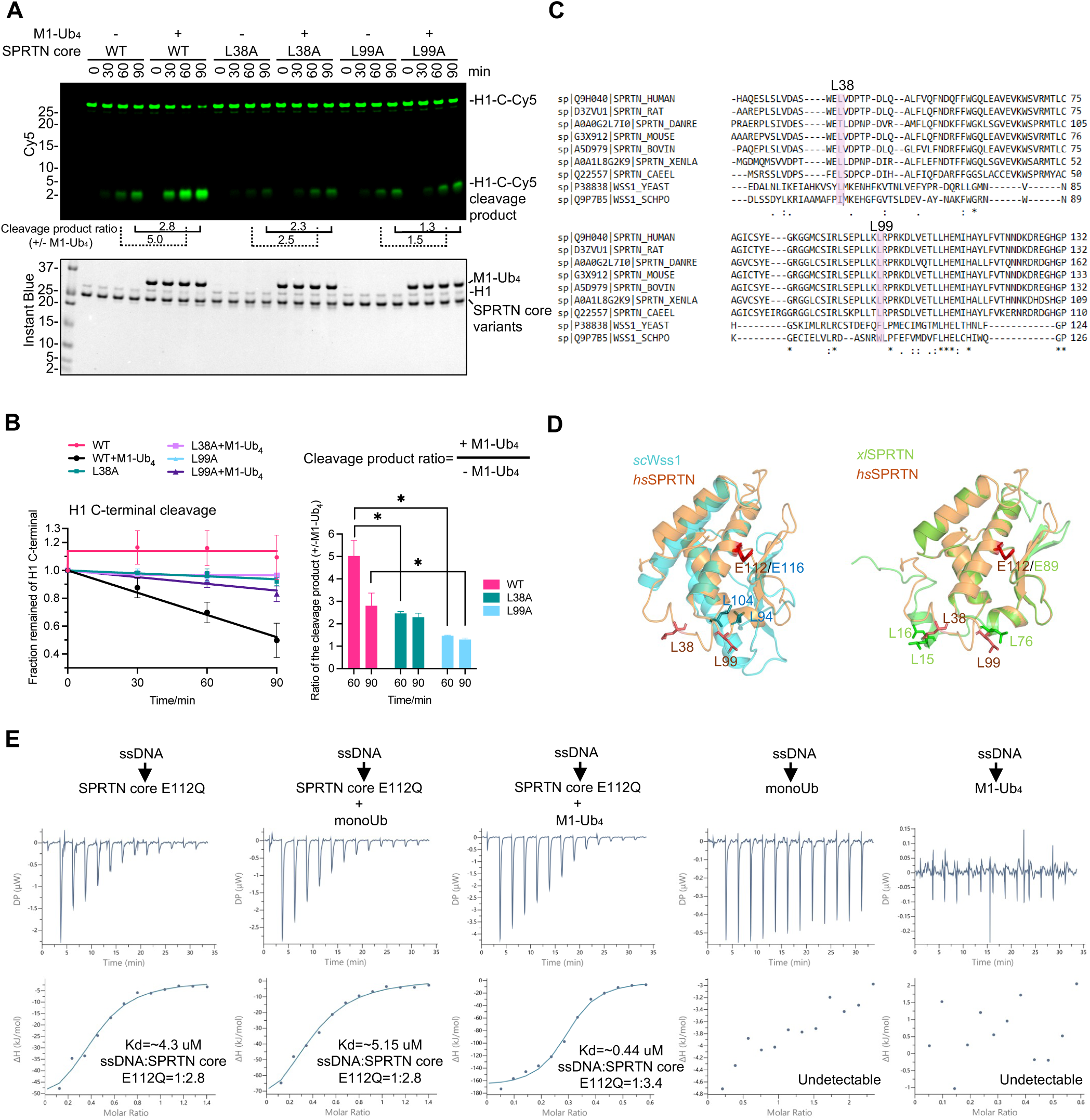
The SPRTN core region of SPRTN contains an intrinsic ubiquitin-binding interface. **(A)** Validation of SPRTN core mutants on the effect of Ub-mediated activation. Recombinant SPRTN core variants (2 µM) and H1-C-Cy5 (1 µM) were incubated in the absence or presence of (M1-Ub_4_, 2 µM) in combination with dsDNA_20/23nt (2.7 µM) with the indicated time at 30°C. The reaction was analysed with SDS-PAGE followed by Cy5-scanning on Typhoon FLA 9500 (GE Healthcare) and Instant Blue staining. Representative figure from 3 repeats. The ratio of cleavage product intensity with/without M1-Ub_4_ at 60 and 90 min for each variant is indicated below the gel. **(B)** Kinetics of the full-length H1-C-Cy5 substrate (C-terminal cleavage rate) and the ratio of cleavage product intensity with/without M1-Ub_4_ at 30 and 90 min, respectively, from Figure 5A. Significant analysis was performed by paired t-test (Prism). n=3. Error bar, SD. *p <0.05. **(C)** Multiple protein sequence alignments of representative SPRTN from vertebrates and Wss1 from yeast by CLUSTAL. **(D)** Superimposition of the human SPRTN MPD domain (AlphaFold Protein Structure Database Entry: AF-Q9H040-F1-v4) with the WLM domain from Wss1 (AlphaFold Protein Structure Database Entry: AF-P38838-F1-v4) (left) or MPD domain from Xenopus SPRTN (AlphaFold Protein Structure Database Entry: AF-A0A1L8G2K9-F1-v4) (right). WLM, Wss1 like metalloprotease domain. **(E)** Isothermal titration calorimetry (ITC) analysis of the binding between ssDNA (20 nt) and SPRTN core (E112Q) in the presence of Ubs (monoUb, M1-Ub_4_, at the equal molar of SPRTN core (E112Q)). The dissociation constant (Kd) and stoichiometry of binding (N) are indicated here and summarized in Table S2.

Sequence alignment also indicated that residues L38 and L99 are highly conserved in vertebrates (Figure 5C). Therefore, we named this Ub binding interface the **<u>U</u>**biquitin-binding interface of **<u>S</u>**prT **<u>D</u>**omain (**USD**). USD of human SPRTN could be very well superimposed with the equivalent leucine residues in *Xenopus* SPRTN protein (Figure 5D). However, the functional homolog of SPRTN in yeast, Wss1, and its WLM protease domain could only partially aligned with the vertebrate/human USD interface. The region in Wss1 corresponding to the human SprT loop containing L38 is absent (Figure 5D). Altogether, our results demonstrate the identification of a Ub-binding interface, the USD, in the SPRTN core region as the essential module for the rapid activation of SPRTN protease in vertebrates.

As the SPRTN core carries both DNA-binding and Ub-binding properties, we used a simplified model to address the ternary corporation consisting of DNA, ubiquitin and SPRTN core. We investigated the binding stoichiometry of ssDNA:SPRTN core in the presence of monoUb or M1-Ub_4_ by ITC (Figures 5E, Table S2). Notably, the binding stoichiometry of ssDNA:SPRTN core changed from ∼1:2.8 without Ub or with monoUb, to ∼1:3.4 in the presence of equal molar of M1-Ub_4_ and SPRTN core. The addition of an equimolar of M1-Ub_4_ and SPRTN core also increased the affinity of SPRTN to ssDNA roughly 10-fold (Figure 5E, Table S2), indicating that the longer Ub chains facilitate the recruitment of SPRTN to DNA.

### Polyubiquitinated substrate rapidly stimulates the SPRTN proteolysis

DPCs are heavily polyubiquitinated, and SPRTN is an essential enzyme for DPC proteolysis repair.^35,39,45^ Thus, we speculated whether ubiquitinated substrates also stimulate SPRTN protease activity.

To this end, we employed several histone H1 substrates where monoUb, M1-Ub_2,_ M1-Ub_4_ or ISG15 were fused to the N-terminus of Cy5-labelled Histone H1 (Cy5-N-H1) for SPRTN proteolysis (Figure 6A). Firstly, SPRTN cleavage kinetics were monitored for the unmodified Cy5-N-H1 and M1-Ub_4_-fused H1 (M1-Ub_4_-Cy5-N-H1). Surprisingly, SPRTN cleavage towards M1-Ub_4_-Cy5-N-H1 was enormously accelerated (less than 2.5 min) (Figures 6B and 6C) compared to the simple addition of free M1-Ub_4_ chains in the reaction (more than 60 min; Figures 2E and 2F). The linear fitting for the whole time course from Cy5-N-H1 and the first 2.5 min from M1-Ub_4_-Cy5-N-H1 was compared at a difference of ∼ 110-fold change, suggesting that Ub chains attached to the substrate stimulate SPRTN proteolytic activity more potently than free Ub chains (Figure 6B and 6C). However, this effect was not so pronounced on SPRTN auto-cleavage (Figures 6B and 6C; bottom panel; ∼ 2.4 fold change; data from the first 5 min for both conditions was linearly fitted). This result was also achieved with the SPRTN core region (26-240 aa; this truncated version of SPRTN does not have a pronounced autocleavage effect) when analysed for histone H1 cleavage (Figures S5A and S5B). Moreover, the rapid cleavage of Cy5-Ub_4_-N-H1 by the SPRTN core region was compromised by the L38A and L99A mutations, albeit the L38A exhibited a milder phenotype (Figures S5C and S5D).

**Figure 6.**
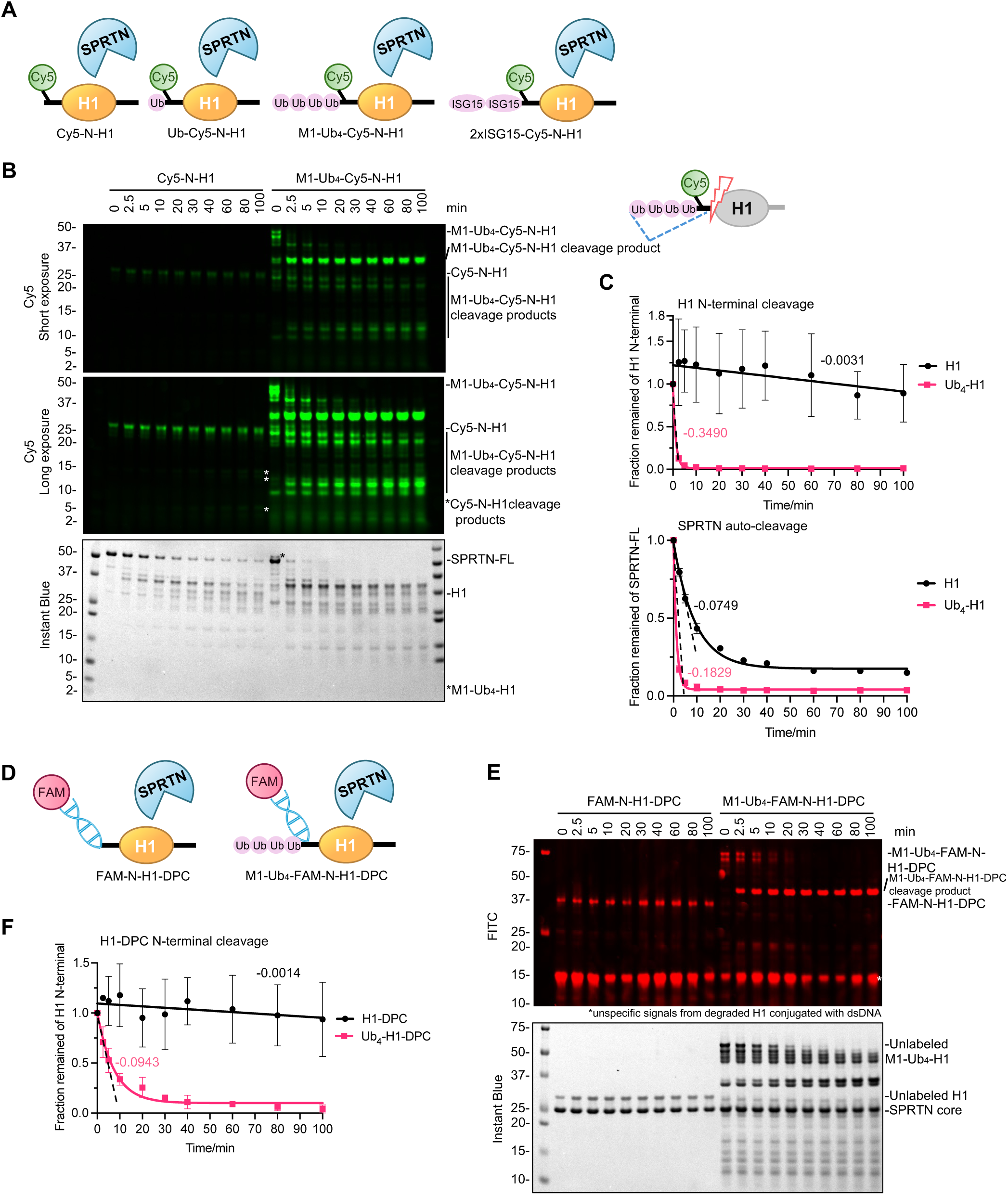
SPRTN rapidly resolves polyubiquitinated DPCs. **(A)** Design of the defined H1 substrates with post-translational modifications for SPRTN cleavage assay (Cy5-labelled all on H1 N-terminal). All the constructs have a His-tag on the N-terminal. **(B)** SPRTN cleavage assay towards Cy5-N-H1 and M1-Ub_4_-Cy5-N-H1. Recombinant full-length SPRTN (2 µM) and H1 substrates (1 µM) were incubated in the presence of dsDNA_20/23nt (2.7 µM) with the indicated time at 30°C. The reactions were analysed by SDS-PAGE followed by Cy5-scanning on an iBright 1500 imaging system (Invitrogen) and Instant Blue staining. Representative figure from 2 repeats. **(C)** Kinetics of the full-length H1 substrates (N-terminal cleavage rate) and the full-length SPRTN (auto-cleavage rate) from Figure 6B. Cy5 and SPRTN-FL signals were analysed by the iBright Analysis Software (Invitrogen). For H1 N-terminal cleavage, the kinetic data for H1 were fitted with simple linear regression (Prism), while the first 2.5 min data points for Ub_4_-H1 were fitted with simple linear regression (Prism). The kinetic data for the whole time course of Ub_4_-H1 were fitted with one phase exponential decay - least squares fit (Prism). For SPRTN auto-cleavage, the kinetic data for the whole time course were fitted with one phase exponential decay - least squares fit (Prism). The kinetic data for the first 5 min were fitted with simple linear regression (Prism). All the slope values for the linear fitting are indicated next to the corresponding dash lines. N=2. Error bar, SD. **(D)** Design of the FAM-dsDNA labelled H1-DPC substrates (FAM-N-H1-DPC and M1-Ub_4_-FAM-N-DPC) for SPRTN cleavage assay. **(E)** SPRTN core cleavage assay towards the defined H1-DPC substrates from Figure 6D. Recombinant SPRTN core (2 µM) and H1-DPC substrates (∼0.1 µM) were incubated at the indicated time at 30°C. The reaction was analysed by SDS-PAGE followed by FITC-scanning on an iBright 1500 imaging system (Invitrogen) and Instant Blue staining. Representative figure from 3 repeats. **(F)** Kinetics of the full-length H1-DPC substrates (N-terminal cleavage rate) from Figure 6E. FITC signals were analysed by the iBright Analysis Software (Invitrogen). Kinetic data for H1-DPC were fitted with simple linear regression (Prism) with the slope value indicated. Kinetic data for Ub_4_-H1-DPC were fitted with one phase exponential decay - least squares fit (Prism). The first 5 min data points from Ub_4_-H1-DPC were fitted with simple linear regression (Prism) with the slope indicated in the dashed line. N=3. Error bar, SD. See also Figure S5.

Notably, a major cleavage product between 25-37 KDa from M1-Ub_4_-Cy5-N-H1 accumulated over time (Figures 6B and S5A). This product retains the N-terminus of H1 (as it was Cy5-visible) and full-length M1-Ub_4_ (as it was detected with an Anti-His antibody) (Figure S5E), indicating that SPRTN did not cleave within the Ub chains. Similar to unmodified H1 (Cy5-N-H1), the cleavage kinetics of mono-ubiquitinated H1 (Ub-Cy5-N-H1) or H1 fused to two tandem molecules of ISG15 (2xISG15-Cy5-N-H1), which mimic the structure of M1-Ub_4_, was not as fast as M1-Ub_4_-Cy5-N-H1 (Figure S5F), in line with our previous observations (see Figure 2E and 2G).

Histone H1 fused with M1-Ub_4_ was conjugated to dsDNA to mimic a modified DPC (M1-Ub_4_-FAM-N-H1-DPC) (Figure 6D). Similar to unconjugated M1-Ub_4_-N-H1, ubiquitinated FAM-N-H1-DPC was a better SPRTN substrate than its non-ubiquitinated version (Figures 6E and 6F). Linear fitting for the whole time course from FAM-N-H1-DPC and the first 5 min from M1-Ub_4_-FAM-N-H1-DPC was compared to better understand the rate difference. It clearly showed that the SPRTN core cleaves the ubiquitinated H1-DPC roughly 67-fold faster than the non-ubiquitinated H1-DPC (Figure 6F). These results collectively indicate that SPRTN proteolysis is greatly activated through polyubiquitinated DPC substrates.

### USD domain binds Ubiquitin chains by avidity, and the UBZ domain by high affinity

Although we have demonstrated that the UBZ domain of SPRTN is not involved in the ubiquitin-activation effect of SPRTN proteolytic activity (Figures 3A and 3B), UBZ still has vital roles in regulating SPRTN by its Ub-binding ability and substrate recruitment. ^35,42–44^ As both the USD and the UBZ domains interact with Ub, we asked if the two domains compete for ubiquitin binding.

To this end, we probed the affinity between the UBZ domain (452-489 aa) and monoUb, M1-Ub_2_ and M1-Ub_4_. ITC clearly showed a similar affinity of UBZ, but not its Ub-binding deficient variant UBZ* (C456AC459A), to monoUb or Ub chains at ∼ 1.6 µM (Figures 7A, S6A and Table S3), which was much higher than the affinity between SPRTN core region and monoUb or Ub chains (Figures 4A and S4A). We also tested UBZ and UBZ* domains by size-exclusion chromatography with multi-angle light scattering (SEC-MALS), which confirmed that they stay in a monomer state (Figure S6B and Table S4). The similar affinity of UBZ to monoUb and Ub chains and the monomeric state suggested that the UBZ domain does not bind to Ub in an avidity manner. Interestingly, according to the AlphaFold prediction of the interaction between the UBZ domain and monoUb, UBZ also interacts mainly with the ubiquitin I44 patch (Figure S6C). To further confirm this, the perturbed surface on ubiquitin upon UBZ titration was also probed by NMR (Figures 7B and S6D). It largely overlapped with the surface identified in the NMR profile of monoUb upon SPRTN core titration (Figure 4C), indicating the possibility that UBZ and SPRTN core (via USD) compete to bind ubiquitin at the same position. To investigate the possible competition effect, the UBZ or UBZ* domain was titrated into a reaction where M1-Ub_4_-Cy5-N-H1 was used as the substrate for SPRTN core proteolytic cleavage (Figure 7C). Interestingly, UBZ, but not UBZ*, only inhibited SPRTN core protease activity when titrated in significant excess (molar ratio UBZ:SPRTN core=25:1) but not in the equimolar ratio (1:1) (Figure 7D). This suggests that if the SPRTN UBZ domain engages with polyubiquitin chains on the substrate, the SPRTN core region can still be accommodated to the same chain. This also aligns with our data where both full-length SPRTN and SPRTN core could rapidly resolve ubiquitinated H1 with similar kinetics (Figure 6B, 6C, S5A and S5B).

**Figure 7.**
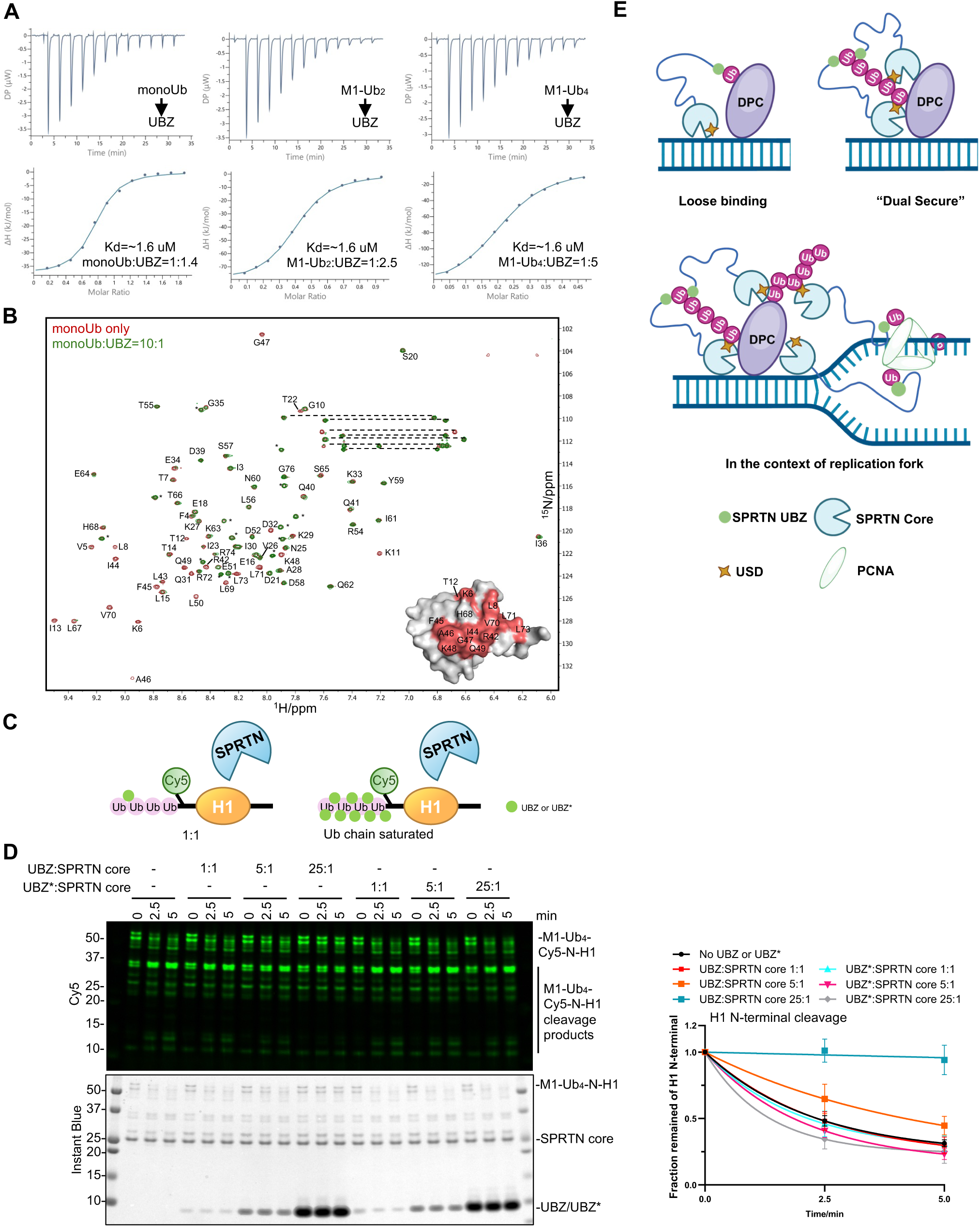
SPRTN UBZ and SPRTN core work in concert to resolve polyubiquitinated DPC substrates. **(A)** Isothermal titration calorimetry (ITC) analysis of the binding between SPRTN UBZ domain and Ubs (monoUb, M1-Ub_2_, M1-Ub_4_). The dissociation constant (Kd) and stoichiometry of binding (N) are indicated here and summarized in Table S3. **(B)** Superimposition of the 950 MHz ^1^H-^15^N HSQC spectra of monoUb alone (in red contours) or with the addition of SPRTN UBZ domain at a molar ratio of 10:1 (in green contours). Spectra were plotted using the MestReNova software. Residues with complete broadening are highlighted in salmon on the monoUb surface (PDB: 1UBQ). The signals were assigned according to previously reported chemical shifts of ubiquitin (BMRB entry: 17769). *Signals from the N-terminal His tag on monoUb. **(C)** Design of the Ub-binding competition between SPRTN core and SPRTN UBZ domain. **(D)** Ub-binding competition between SPRTN core and SPRTN UBZ or UBZ* (C456AC459A). Recombinant SPRTN core (2 µM) and M1-Ub_4_-Cy5-N-H1 (1 µM) were incubated in the presence of dsDNA_20/23nt (2.7 µM) in difference ratio of UBZ:SPRTN core from 1:1 to 25:1 with the indicated time at 30°C. The reaction was analysed with SDS-PAGE followed by Cy5-scanning on an iBright 1500 imaging system (Invitrogen) and Instant Blue staining. Right panel: Kinetics of the full-length M1-Ub_4_-Cy5-N-H1 (N-terminal cleavage rate). Kinetic data were fitted with one phase exponential decay - least squares fit (Prism). N=3. Error bar, SD. **(E)** Models of SPRTN UBZ and SPRTN core working in concert to resolve ubiquitinated DPC substrates. See also Figure S6.

The higher affinity between UBZ and Ub (1.6 µM) supports the idea that UBZ is the general Ub sensor. On the other hand, the avidity effect between the SPRTN core and Ub chains (from ∼770 µM for monoUb to ∼35 µM for M1-Ub_6_) and the ability of longer ubiquitin chains to efficiently stimulate SPRTN protease activity suggest that ubiquitin chain is another factor in bringing SPRTN close to the DPC substrates, in addition to DNA. These results demonstrate that the N-terminally located SPRTN core (with its USD interface) and the C-terminally located UBZ domain work together rather than compete to resolve polyubiquitinated substrates/DPCs.

## DISCUSSION

Replication-coupled DPC proteolysis repair is an essential DNA repair mechanism for genome stability and protection from accelerated ageing and cancer.^28^ SPRTN protease emerged as the crucial enzyme for this repair, but how its specificity is achieved towards DPCs rather than the replisome is still enigmatic. Interestingly, DPC and DPC-like proteins are heavily polyubiquitinated by various E3-ubiquitin ligases, especially in response to chemotherapy drugs or formaldehyde,^8,27,35,45,46^ but whether and how polyubiquitinated substrates/DPCs regulate SPRTN proteolysis has not yet been elucidated.

The work presented here demonstrates that SPRTN is a specialised protease localised to and rapidly activated by polyubiquitinated chains on the substrates/DPCs using its intrinsic ubiquitin-binding modules, UBZ and USD, respectively. SPRTN cleaves the polyubiquitinated substrates irrespective of the chain linkages, making SPRTN a universal DPC protease for polyubiquitinated DPCs that can occur regardless of the E3-ubiquitin ligase involved in DPC repair. This work clarifies the essential question in the field: How does the SPRTN protease specifically process DPCs and thus avoid the cleavage of other replicative proteins such as PCNA?

### Polyubiqutination of DPCs governs SPRTN’s specificity and its rapid proteolysis

SPRTN was identified as a protein that contains a Zn-finger Ub-binding (UBZ) domain that regulates PCNA mono-ubiquitination status during translesion DNA synthesis and a cofactor of the Ub-dependent ATPase p97, the central component of the Ub system.^42–44,47,48^ However, how Ub or the ubiquitination of substrates/DPCs regulates SPRTN’s protease activity and specificity is enigmatic. There is a consensus in the field that the UBZ domain, by binding to monoUb or polyUb chains, regulates SPRTN stability and its recruitment to DPCs, respectively.^31,39^ Still, it does not restrict its enzymatic activity or substrate specificity.

It is evident that many factors stimulate SPRTN proteolysis and cellular response to DPC-inducing agents^20,24,25,43,44,47–52^, but how the polyubiquitination of substrates/DPCs, the robust posttranslational signal on DPCs induced by formaldehyde or chemotherapy drugs treatment,^8,35,39,45^ regulates SPRTN proteolysis is not known. One model is that when the replicative helicase CMG approaches DPCs, it bypasses the DPC lesion but is uncoupled from the PCNA/DNA polymerase complex, thus forming a single-strand DNA structure where DPCs are attached.^32^ When the PCNA/DNA polymerase complex approaches the DPC, this is a perfect DNA structure (ss/dsDNA junction) for SPRTN activation.^3^ However, even though this ss/dsDNA junction is indeed the best DNA substrate for SPRTN proteolysis, and we confirm it too (Figure 1), it does not explain how SPRTN gains specificity between DPCs and replication machinery. Moreover, this ss/dsDNA structure also takes hours to activate SPRTN proteolysis *in vitro*.^3,19,30^

Specifically, we identified that DPC polyubiquitination is the primary signal for SPRTN’s specificity and rapid proteolysis of DPCs. SPRTN is an Ub-activated protease with a previously unidentified Ub-binding interface (USD) in its catalytic-core domain. USD barely detects monoUb (affinity for mono Ub is ∼770 µM, Figure S4C) but rapidly increases its affinity in an avidity mode for the growing ubiquitin chains (polyubiquitination; USD affinity to M1-Ub_4_ is around 70 µM, and further increased to around 35 µM with M1-Ub_6_, Figures 4A, S4A and Table S1). Together with the DNA-binding ability from the ZBD and BR domain, this Ub-threshold property from USD largely defines the substrate selectivity and the timely enzymatic efficiency of SPRTN. In contrast to USD, the C-terminal UBZ domain has a strong affinity for either monoUb or Ub chains (around 1.6 µM, Figure 7A and Table S3), but won’t outcompete USD when binding to longer ubiquitin chains. Based on our results, we propose a model where two intrinsic Ub-binding modules of SPRTN, USD and UBZ regulate the spatiotemporal and rapid proteolysis of polyubiquitinated DPCs in a “dual secure” mode (Figure 7E).

### A model of SPRTN specificity at/around the DNA replication fork

SPRTN was identified to bind PCNA over its PIP box and is associated with DNA replicative polymerases.^25,42–44,48,53^ Uncontrolled SPRTN protease activity at/around the DNA replication fork can be detrimental to cells, as it has been believed that SPRTN is an unspecific protease. Our model suggests that SPRTN can be equally recruited to monoubiquitinated PCNA via its PIP box and UBZ domain. Binding data from SPR also support that the SPRTN PIP-UBZ module has a decent affinity to PCNA and monoUb-PCNA (Figures S6E and Table S5). As PCNA is a homotrimer and each homotrimer contains one mono-ubiquitinated site at Lysine 164, it is reasonable to speculate that one molecule of monoUb-PCNA homotrimer can accommodate three molecules of SPRTN by binding to both the PIP box and UBZ domain (Figure 7E). Once the SPRTN C-terminal tail is stabilised on PCNA by the PIP box and UBZ domain, USD allows the recruitment of SPRTN protease core to the sites of polyubiquitinated DPCs either during the ongoing DNA replication (non-ubiquitinated PCNA) or during stalled DNA replication fork (monoUb-PCNA) (Figure 7E). If the DNA replication fork approaches DPCs, and the DPCs are heavily polyubiquitinated, SPRTN rapidly increases its affinity and specificity towards the growing ubiquitin chains and, consequently, DPCs.

On the other hand, the Ub-threshold of USD could avoid the degradation of monoUb-PCNA or DNA polymerases nearby as these proteins are not heavily polyubiquitinated during unperturbed DNA replication fork progression.^54–59^ As SPRTN possesses a large flexible region between the N-terminal USD-containing protease domain and the C-terminal UBZ domain, indicated by AlphaFold prediction (Figure S6F, AlphaFold Protein Structure Database Entry: AF-Q9H040-F1-v4), this also suggests that the catalytic domain of SPRTN could reach polyubiquitinated substrates/DPCs around PCNA/DNA replication while still being attached to PCNA. Our model indicates that SPRTN facilitates the rapid proteolysis of polyubiquitinated substrates/DPCs with high selectivity on the replication fork.

### SPRTN vs 26S Proteasome in the processing of polyubiquitinated DPCs

The K48-linked poly-ubiquitination of proteins is the main ubiquitin signal that governs substrate degradation by the 26S proteasome.^60^ On the contrary, SPRTN protease was previously identified to cleave its substrates in a ubiquitination-independent manner *in vitro*.^3,19,22,25^ Based on these facts, there is a general opinion in the field that the 26S proteasome processes polyubiquitinated DPCs, and SPRTN processes those not ubiquitinated.^27,61^ However, it’s puzzling that SPRTN proteolysis on non-ubiquitinated DPCs is still dependent on the UBZ domain, and it was suggested that SPRTN is recruited by a ubiquitinated protein around DPC but other than the DPC itself.^27^

Our work indicates that the polyUb signal on the substrates/DPCs is the main catalyst for SPRTN proteolysis. Without the polyubiquitination of substrates/DPCs, SPRTN is an inefficient enzyme that takes hours to cleave its substrates, which contradicts its role in the rapid resolution of DPCs during DNA synthesis. By recognizing the polyUb chains attached to substrates/DPCs, SPRTN becomes a rapid and processive protease that cleaves its substrates within a few minutes (2.5-10 min, Figure 6). As SPRTN activation by polyubiquitin chains is not linkage-specific, contrary to the proteasome, we propose that SPRTN is a universal protease for removing ubiquitinated DPCs. The polyubiquitin chains on DPCs serve as an excellent fine-tuning signal to regulate SPRTN selectivity and proteolysis towards DPCs, thus avoiding unspecific cleavage of non-ubiquitinated replication-associated proteins during DNA synthesis.^54,55^

### Ethical consent

Not applicable.

## RESOURCE AVAILABILITY

### Lead contact

Further information and requests for resources and reagents should be directed to and will be fulfilled upon reasonable request by the Lead Contact, Kristijan Ramadan.

### Materials Availability

All plasmids are available on request.

### Data and Code Availability

This study did not generate code or reposited datasets.

## ACKNOWLEDGMENTS

The authors thank Dr Benjamin Stieglitz for providing the Ub, NEDD8 and mUbe1 constructs and Dr Gemma Harris from the UKRI Research Complex at Harwell for performing SEC-MALS experiments and analysis.

This work received support from the Medical Research Council programme (Grant Ref. MR/X006409/1), Breast Cancer Now (Grant Ref. 2022.11PR1570), Ministry of Education-Strat-Up Grant, Singapore (023917-00001), Toh Kian Chui Distinguished Professorship Award/LKC Medicine, Singapore to K.R. The 950 MHz spectrometer was upgraded with funding from the University of Oxford Wellcome Institutional Strategic Support Fund, the John Fell Fund, the Edward Penley Abraham Cephalosporin Fund, and the Engineering and Physical Sciences Research Council (Grant Ref: EP/R029849/1). J.A.N. acknowledges support for his research from the following grants NCI P01 CA092584 and CRUK A24759.

## AUTHOR CONTRIBUITIONS

Conceptualisation, W.S. and K.R.; Methodology, Formal Analysis and Investigation, W.S., J.A.N., C.R., Y.Z.; Validation, W.S., J.A.N.; Resources, W.S., P.E., X.Z., A.C., R.R., P.M.; Data Curation, all authors; Writing – Original Draft, W.S. and K.R.; Writing – Review & Editing, W.S., A.R. and K.R. with input from all authors; Visualisation, W.S. Y.Z.; Supervision, K.R.; Funding Acquisition, K.R.

## Supplemental Figures and Tables

**Supplemental Figure 1.**
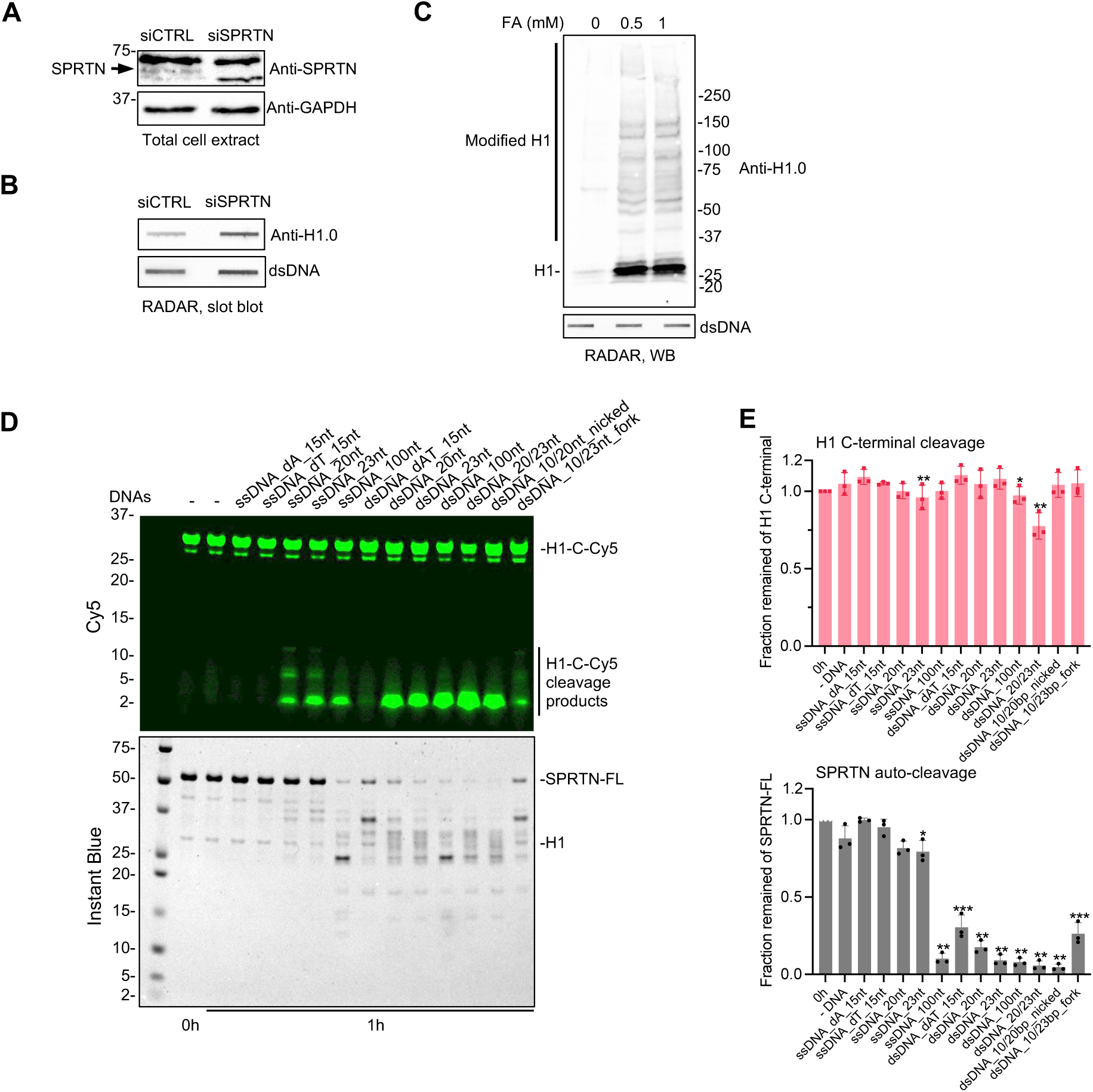
SPRTN proteolysis is activated by both ssDNA and dsDNA. **(A)** Confirmation of SPRTN depletion in U2OS cell line by western blot. **(B)** Total DPCs were isolated from the SPRTN-depleted cells (U2OS cell line) by RADAR and analysed by slot blot. **(C)** Total DPCs were isolated from the formaldehyde-treated cells (Hek293 cell line) by RADAR. Total DPCs were analysed by SDS-PAGE followed by western blot. **(D)** DNA screening for SPRTN cleavage assay. Recombinant full-length SPRTN (2 µM) and H1-C-Cy5 (1 µM) were incubated in the absence or presence of indicated DNA for 1h at 30°C. The reaction was analysed by SDS-PAGE followed by Cy5-scanning on Typhoon FLA 9500 (GE Healthcare) and Instant Blue staining. Representative figure from 3 repeats. **(E)** Quantification of the signal from the full-length H1-C-Cy5 substrate (C-terminal cleavage rate) and the full-length SPRTN (auto-cleavage rate) from Figure S1D. Cy5 signals were analysed by ImageJ. SPRTN-FL signals were analysed by the iBright Analysis Software (Invitrogen). Significant analysis was performed by comparing each data point to “-DNA” by paired t-test (Prism) (Data from 0h time point was excluded from analysis). n=3. Error bar, SD. *p <0.05; **p <0.005; ***p <0.0005.

**Supplemental Figure 2.**
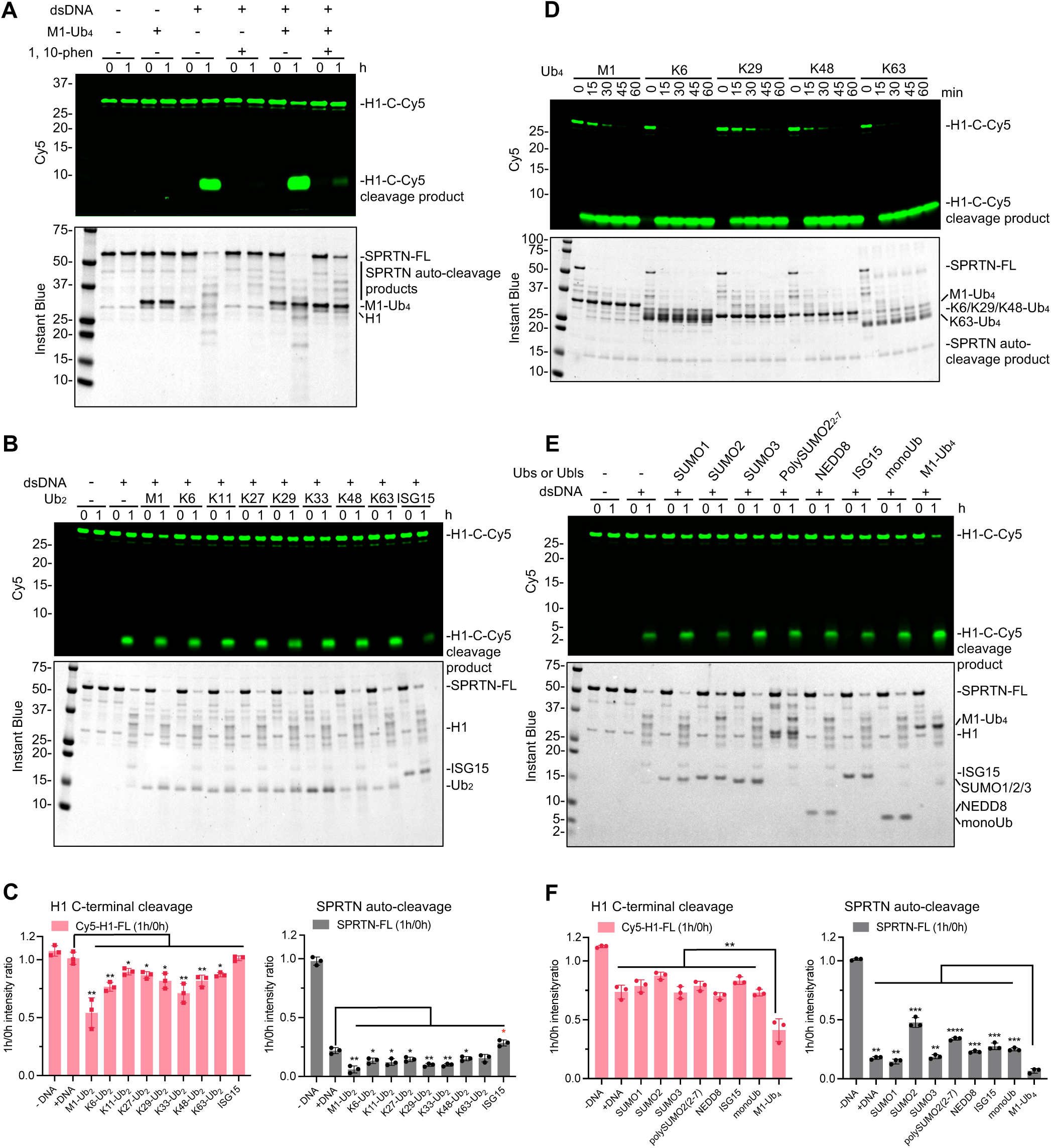
Ubs but not Ub-like proteins activate SPRTN proteolysis. **(A)** SPRTN cleavage assay towards H1-C-Cy5 in the combination of dsDNA_20/23nt with M1-Ub_4_ and 1,10-phenanthroline (1,10-phen). Recombinant full-length SPRTN (2 µM) and H1-C-Cy5 (1 µM) were incubated with different combination of dsDNA_20/23nt (2.7 µM), M1-Ub_4_ (2 µM) and 1,10-phenanthroline (0.5 mM) for 1h at 30°C. Representative figure from 3 repeats. **(B)** Ub_2_ screening for SPRTN cleavage assay. Recombinant full-length SPRTN (2 µM) and H1-C-Cy5 (1 µM) were incubated with indicated Ub_2_ or ISG15 (2 µM) in the presence of dsDNA_20/23nt (2.7 µM) for 1h at 30°C. Representative figure from 3 repeats. **(C)** Quantification of the signal from the full-length H1-C-Cy5 substrate (C-terminal cleavage rate) and the full-length SPRTN (auto-cleavage rate) from Figure S2B. n=3. Error bar, SD. **(D)** Ub_4_ screening for SPRTN cleavage assay. Recombinant full-length SPRTN (2 µM) and H1-C-Cy5 (1 µM) were incubated with indicated Ub_4_ (2 µM) in the presence of dsDNA_20/23nt (2.7 µM) with indicated time at 30°C. Representative figure from 3 repeats. **(E)** Ubl screening for SPRTN cleavage assay. Recombinant full-length SPRTN (2 µM) and H1-C-Cy5 (1 µM) were incubated with indicated Ubs or Ubls (2 µM) in the presence of dsDNA_20/23nt (2.7 µM) for 1h at 30°C. Representative figure from 3 repeats. **(F)** Quantification of the signal from the full-length H1-C-Cy5 substrate (C-terminal cleavage rate) and the full-length SPRTN (auto-cleavage rate) from Figure S2E. All the reactions were analysed by SDS-PAGE followed by Cy5-scanning on Typhoon FLA 9500 (GE Healthcare) and Instant Blue staining. Cy5 signals were analysed by ImageJ. SPRTN-FL were analysed by the iBright Analysis Software (invitrogen). Significant analysis from Figure S2C and S2F was performed by paired t-test (Prism) (Data from “-DNA” was excluded from analysis). *p <0.05; **p <0.005; ***p <0.0005.

**Supplemental Figure 3.**
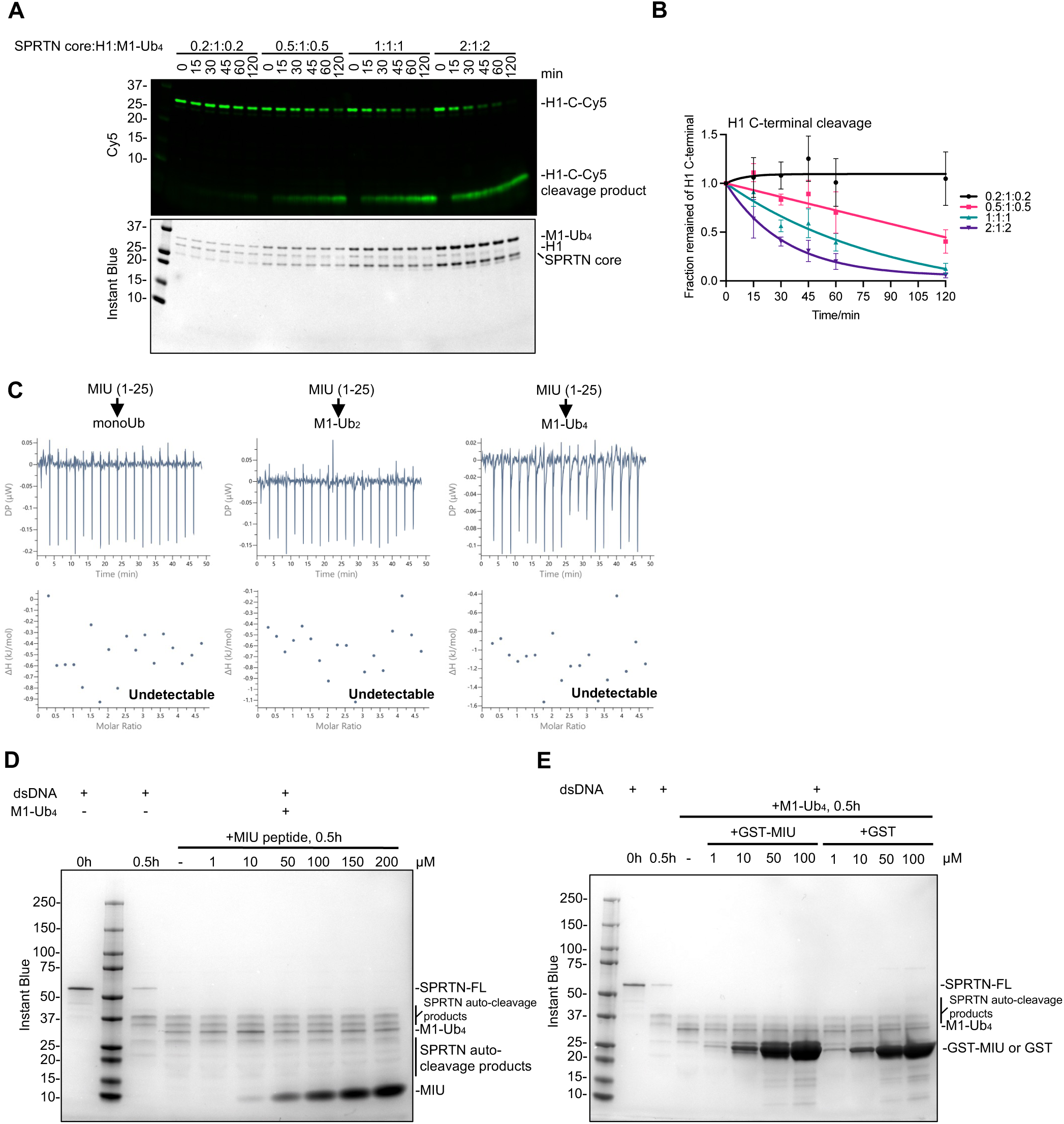
SPRTN core region of SPRTN is sufficient for the rapid activation of SPRTN protease mediated by Ub. **(A)** SPRTN core cleavage assay towards H1-C-Cy5 with M1-Ub_4_. Recombinant SPRTN core and H1-C-Cy5 were incubated with M1-Ub_4_ with the indicated ratio in the presence of dsDNA_20/23nt (2.7 µM) with the indicated time at 30°C. H1-C-Cy5 was kept constantly at 1 µM in all conditions. The reactions were analysed by SDS-PAGE followed by Cy5-scanning on an iBright 1500 imaging system (Invitrogen) and Instant Blue staining. Representative figure from 3 repeats. **(B)** Cleavage kinetics of the signal from the full-length H1-C-Cy5 substrate (C-terminal cleavage rate) from Figure S3A. Cy5 and SPRTN-FL signals were analysed by the iBright Analysis Software (Invitrogen). Kinetic data were fitted with one phase exponential decay - least squares fit (Prism). n=3. Error bar, SD. **(C)** Isothermal titration calorimetry (ITC) analysis of the binding between MIU peptide (1-25 aa) and Ubs (monoUb, M1-Ub_2_, M1-Ub_4_). **(D)** Competition binding assay of MIU peptide to M1-Ub_4_. SPRTN auto-cleavage was monitored. Recombinant full-length SPRTN (2 µM) and M1-Ub_4_ (1 µM) were incubated with MIU peptide with indicated concentration in the presence of dsDNA_20/23nt (2.7 µM) for 0.5h at 30°C. The reaction was analysed by SDS-PAGE. **(E)** Competition binding assay of GST-MIU to M1-Ub_4_. SPRTN auto-cleavage was monitored. Recombinant full-length SPRTN (2 µM) and M1-Ub_4_ (1 µM) were incubated with GST-MIU or GST with indicated concentration in the presence of dsDNA_20/23nt (2.7 µM) for 0.5h at 30°C. The reaction was analysed by SDS-PAGE.

**Supplemental Figure 4.**
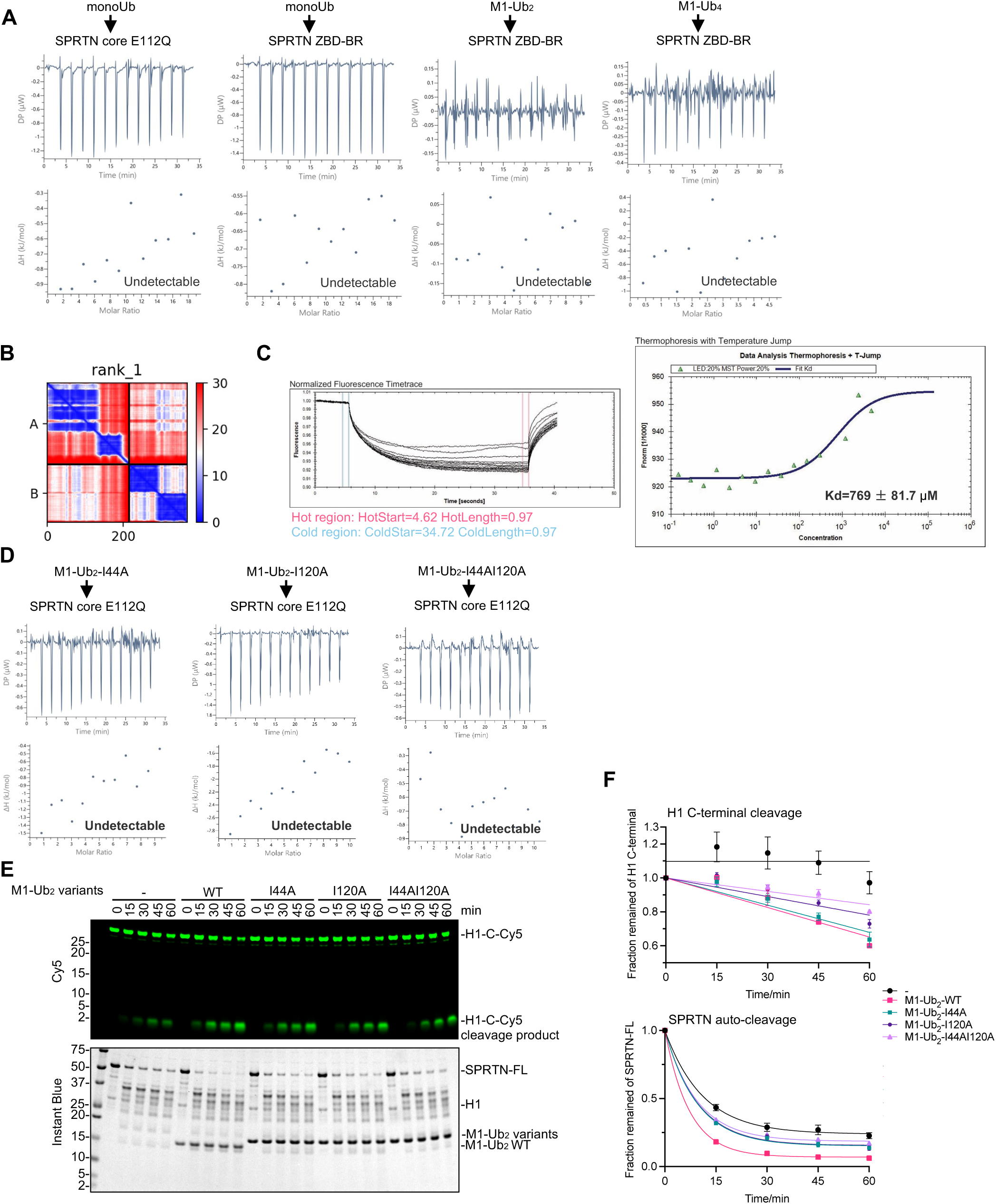
SPRTN core interacts with I44 patch of Ub in an avidity manner. **(A)** Isothermal titration calorimetry (ITC) analysis of the binding between SPRTN core (E112Q) and monoUb, SPRTN ZBD-BR and Ubs (monoUb, M1-Ub_2_, M1-Ub_4_). Details can also be found in Table S1. **(B)** PAE plot of the ColaFold2 prediction of the interaction between SPRTN core and M1-Ub_2_. **(C)** MST analysis of the affinity between SPRTN core (E112Q) and monoUb. The dissociation constant (Kd) is indicated. **(D)** Isothermal titration calorimetry (ITC) analysis of the binding between SPRTN core (E112Q) and M1-Ub_2_ variants (I44A, I120A, I44AI120A). Details can be also found in Table S1. **(E)** Validation of M1-Ub_2_ variants on the effect of SPRTN activation. Recombinant full-length SPRTN (2 µM) and H1-C-Cy5 (1 µM) were incubated in the presence of dsDNA_20/23nt (2.7 µM) in combination with M1-Ub_2_ variants (WT, I44A, I120A, I44AI120A, all at 2 µM) with the indicated time at 30°C. The reaction was analysed with SDS-PAGE followed by Cy5-scanning on Typhoon FLA 9500 (GE Healthcare) and Instant Blue staining. Representative figure from 3 repeats. **(F)** Kinetics of the full-length H1-C-Cy5 substrate (C-terminal cleavage rate) and the full-length SPRTN (auto-cleavage rate) from Figure S4E. Cy5 signals were analysed by ImageJ. SPRTN-FL signals were analysed by the iBright Analysis Software (Invitrogen). Kinetic data were fitted with one phase exponential decay - least squares fit (Prism). N=3. Error bar, SD.

**Supplemental Figure 5.**
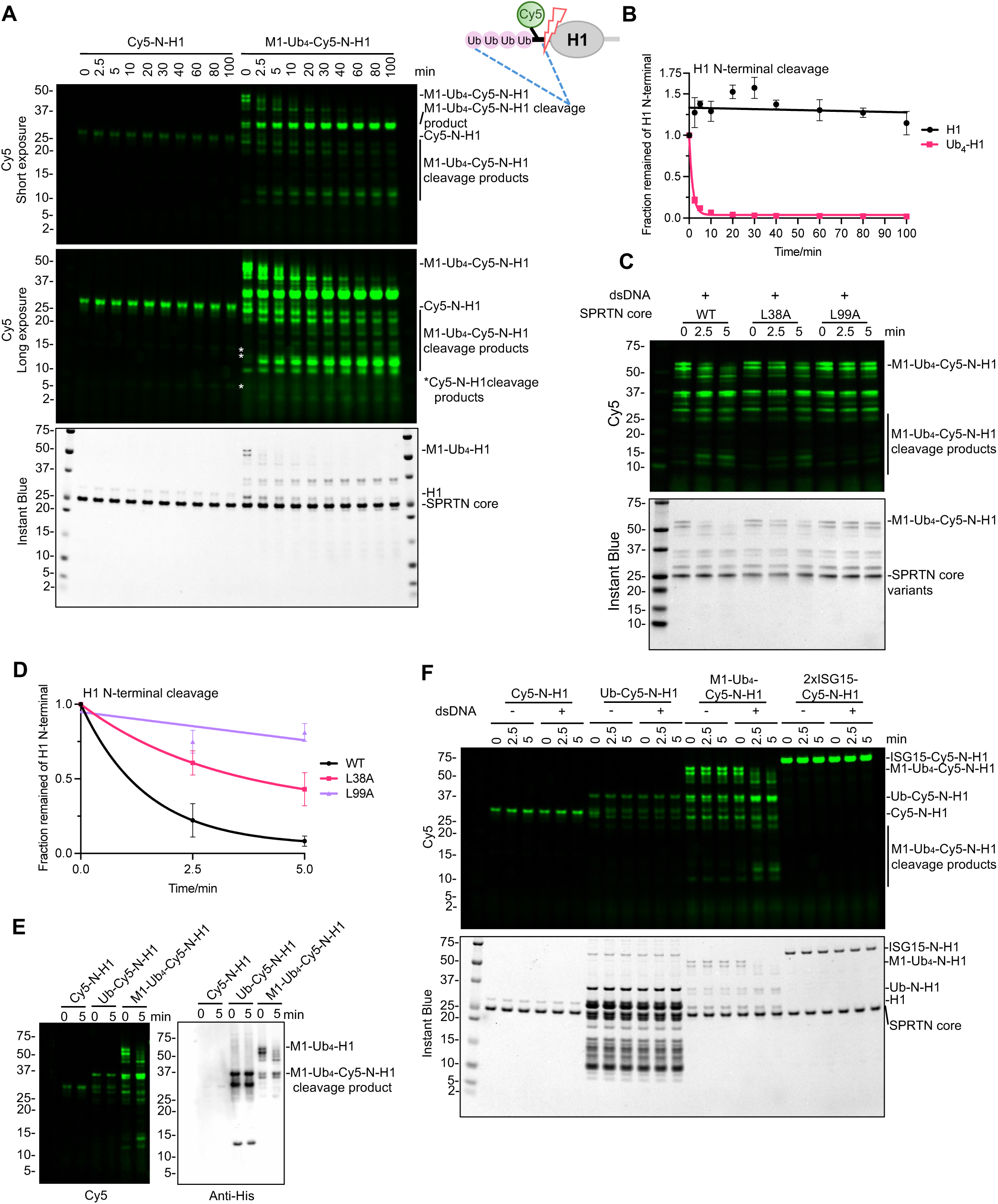
SPRTN core rapidly resolves polyubiquitinated DPCs. **(A)** SPRTN core cleavage assay towards Cy5-N-H1 and M1-Ub_4_-Cy5-N-H1. Recombinant SPRTN core (2 µM) and H1 substrates (1 µM) were incubated in the presence of dsDNA_20/23nt (2.7 µM) with the indicated time at 30°C. Representative figure from 3 repeats. **(B)** Kinetics of the full-length H1 substrates (N-terminal cleavage rate) from Figure S5A. Kinetic data for H1 were fitted with simple linear regression. Kinetic data for Ub_4_-H1 were fitted with one phase exponential decay - least squares fit (Prism). N=3. Error bar, SD. **(C)** Activity of SPRTN core mutants towards M1-Ub_4_-Cy5-N-H1. Recombinant SPRTN core variants (2 µM) and M1-Ub_4_-Cy5-N-H1 (1 µM) were incubated in the presence of dsDNA_20/23Nt (2.7 µM) with the indicated time at 30°C. Representative figure from 3 repeats. **(D)** Kinetics of the full-length M1-Ub_4_-Cy5-N-H1 (N-terminal cleavage rate) from Figure S5E. Kinetic data were fitted with one phase exponential decay - least squares fit (Prism). N=3. Error bar, SD. **(E)** Western blot analysis of the SPRTN core cleavage assay towards the H1 substrates. Recombinant SPRTN core (2 µM) and H1 substrates (1 µM) were incubated in the presence of dsDNA_20/23nt (2.7 µM) with the indicated time at 30°C. The reactions were analysed by SDS-PAGE followed by Cy5-scanning and then applied to western blot by Anti-His antibody. **(F)** SPRTN core cleavage assay towards the defined H1 substrates from Figure 6A. Recombinant SPRTN core (2 µM) and H1 substrates (1 µM) were incubated in the presence of dsDNA_20/23nt (2.7 µM) with the indicated time at 30°C. Representative figure from 3 repeats. All the reactions were analysed by SDS-PAGE followed by Cy5-scanning on an iBright 1500 imaging system (Invitrogen) and Instant Blue staining. Cy5 signals from Figure S5A and S5E were analysed by the iBright Analysis Software (Invitrogen).

**Supplemental Figure 6.**
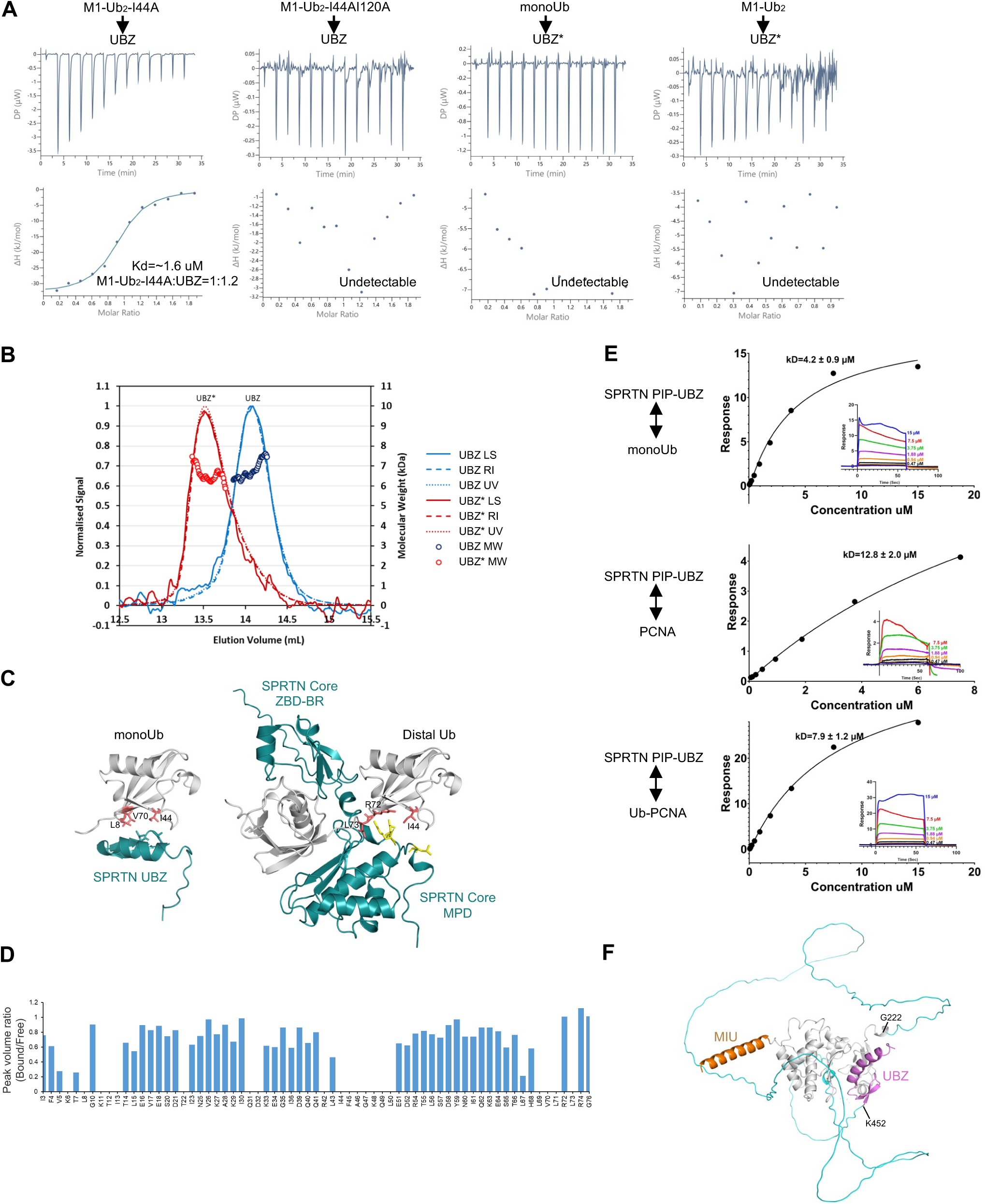
SPRTN UBZ and SPRTN core work in concert to resolve polyubiquitinated DPC substrates. **(A)** Isothermal titration calorimetry (ITC) analysis of the binding between SPRTN UBZ (or UBZ* domain) and Ubs (monoUb, M1-Ub_2_ variants). The dissociation constant (Kd) and stoichiometry of binding (N) are indicated. Details can also be found in Table S3. **(B)** SEC-MALS analysis of SPRTN UBZ and UBZ* domain. Parameters are listed in the Table S4. LS: light scattering; RI: refractive Index. **(C)** Ub-binding comparison between SPRTN UBZ and SPRTN core. Complex models are predicted by AlphaFold3. monoUb and the distal Ub from M1-Ub_2_ are imposed in the same orientation. **(D)** Peak volume ratio of bound/free states for individual residues from Figure 7B. Absolute peak volume was quantified by the MestReNova software. Peaks that are completely broadened upon the addition of UBZ to monoUb have a peak volume ratio of 0. **(E)** Predicted SPRTN-FL structure (AlphaFold Protein Structure Database Entry: AF-Q9H040-F1-v4). The unstructured region (G222-K452) is coloured in cyan. The N-terminal MIU domain is coloured in orange. The C-terminal UBZ domain is colored in magenta. **(F)** Surface plasmon resonance (SPR) analysis of the interaction between SPRTN PIP-UBZ and monoUb/PCNA/monoUb-PCNA. The dissociation constant (Kd) of binding is indicated here and listed in Table S5.

## Supplemental tables

**Table S1^a^:**
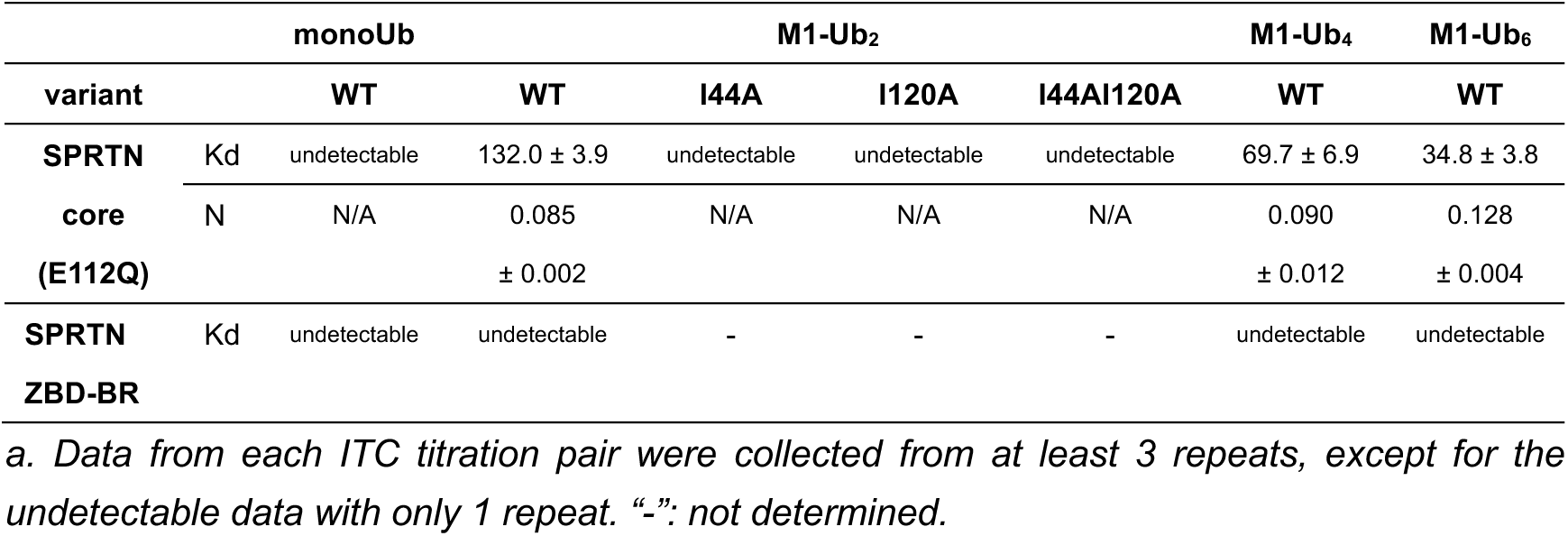
Affinity summary of SPRTN core (E112Q) with ubiquitin from ITC (Unit: µM)

**Table S2^a^:** Affinity summary of ssDNA (20 nt) with SPRTN core (E112Q) in the presence of ubiquitin from ITC (Unit: µM)

|  |  | SPRTN core E112Q | SPRTN core E112Q<br>+ | SPRTN core E112Q<br>+ |
| --- | --- | --- | --- | --- |
|  |  |  | monoUb | M1-Ub<sub>4</sub> |
| **ssDNA** | Kd | 4.33 ± 0.98 | 5.15 ± 1.36 | 0.44 ± 0.03 |
| **(20nt)** | N | 0.36 ± 0.05 | 0.36 ± 0.02 | 0.29 ± 0.01 |

a. Data from each ITC titration pair were collected from at least 3 repeats.

**Table S3^a^:**
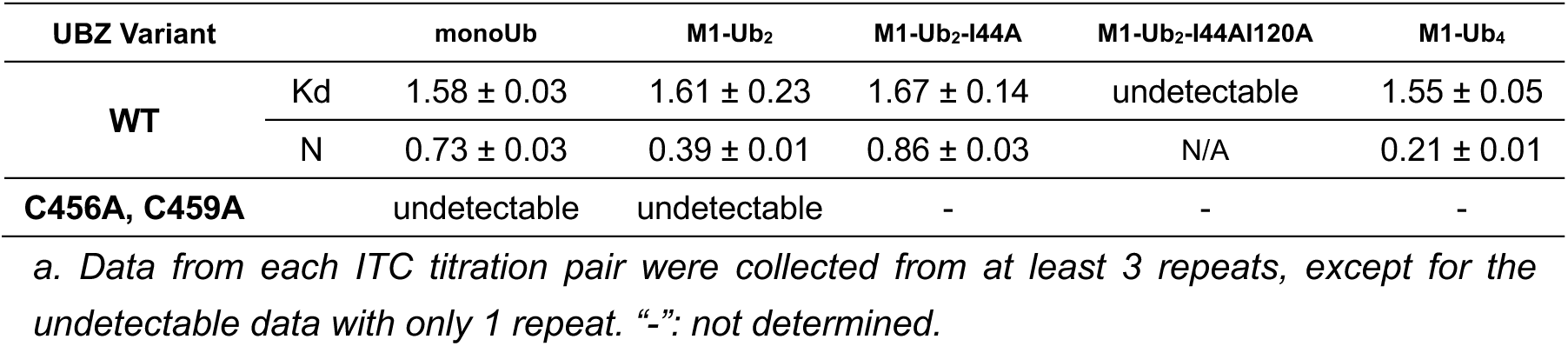
Affinity summary of SPRTN UBZ with ubiquitin from ITC (Unit: µM)

**Table S4:**
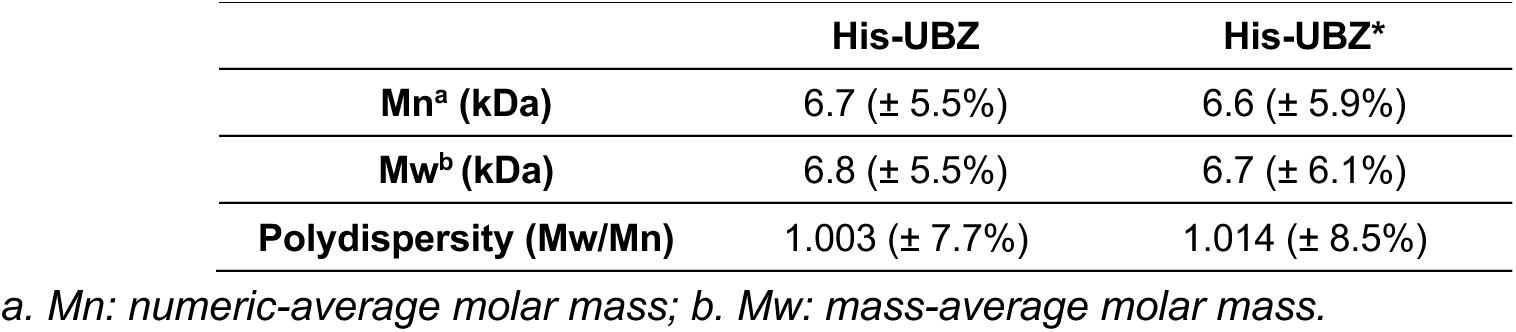
Parameters of SPRTN UBZ and UBZ* from SEC-MALS.

**Table S5:** Affinity summary of SPRTN PIP-UBZ with monoUb/PCNA/Ub-PCNA from SPR (Unit: µM)

|  |  | monoUb | PCNA | Ub-PCNA |
| --- | --- | --- | --- | --- |
| **PIP-UBZ** | Kd | 4.2 ± 0.9 | 12.8 ± 2.0 | 7.9 ± 1.2 |

## STAR METHODS

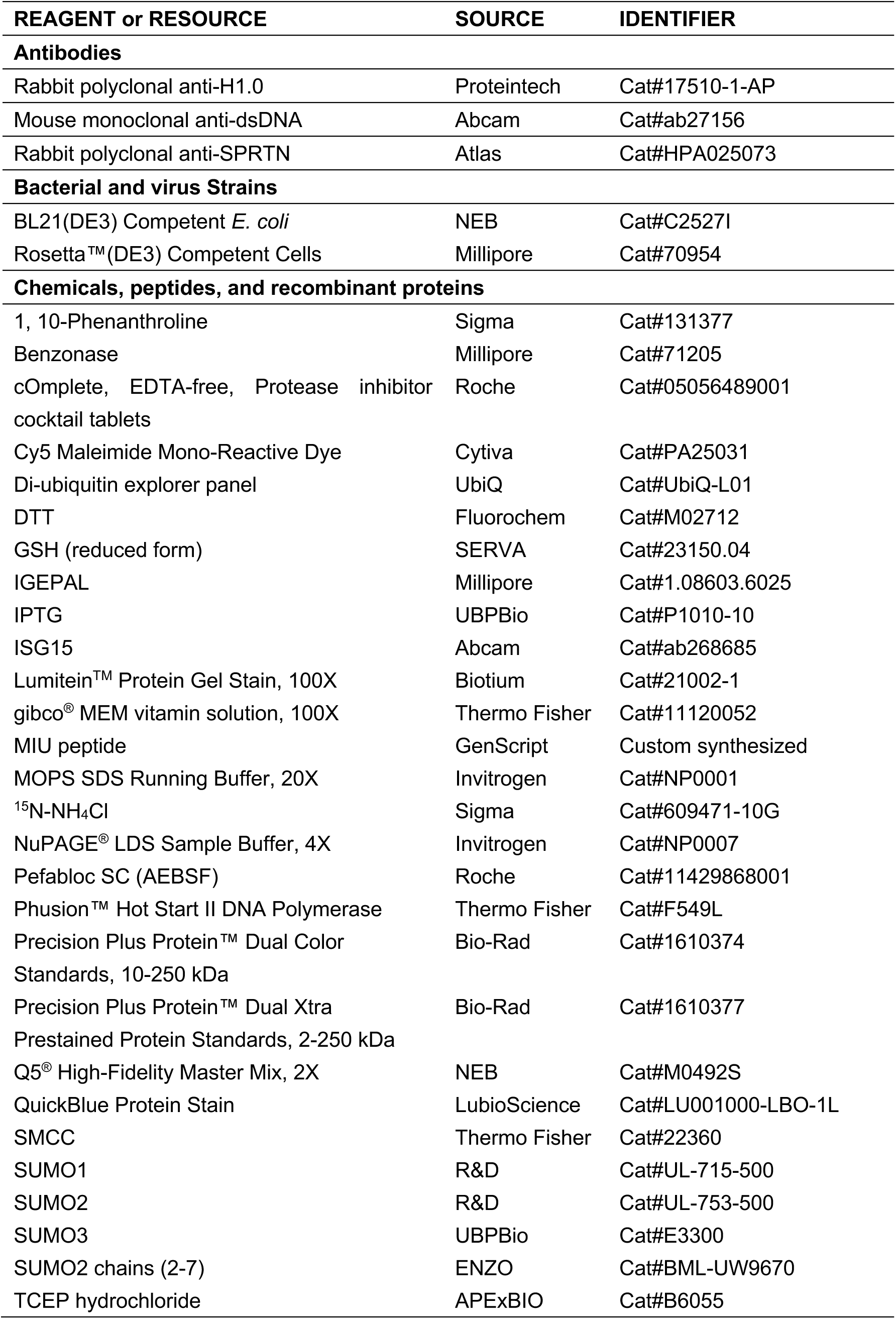
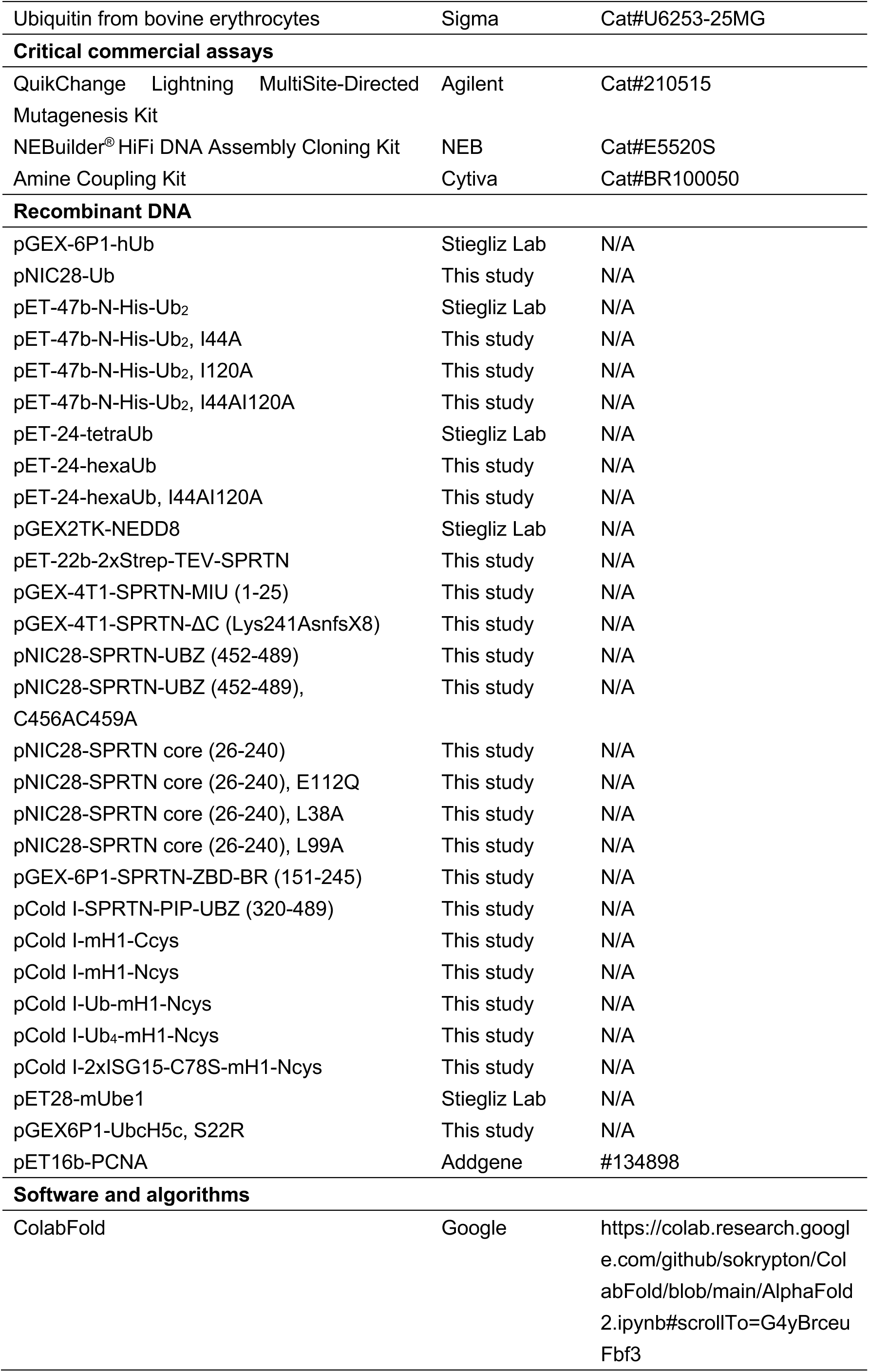
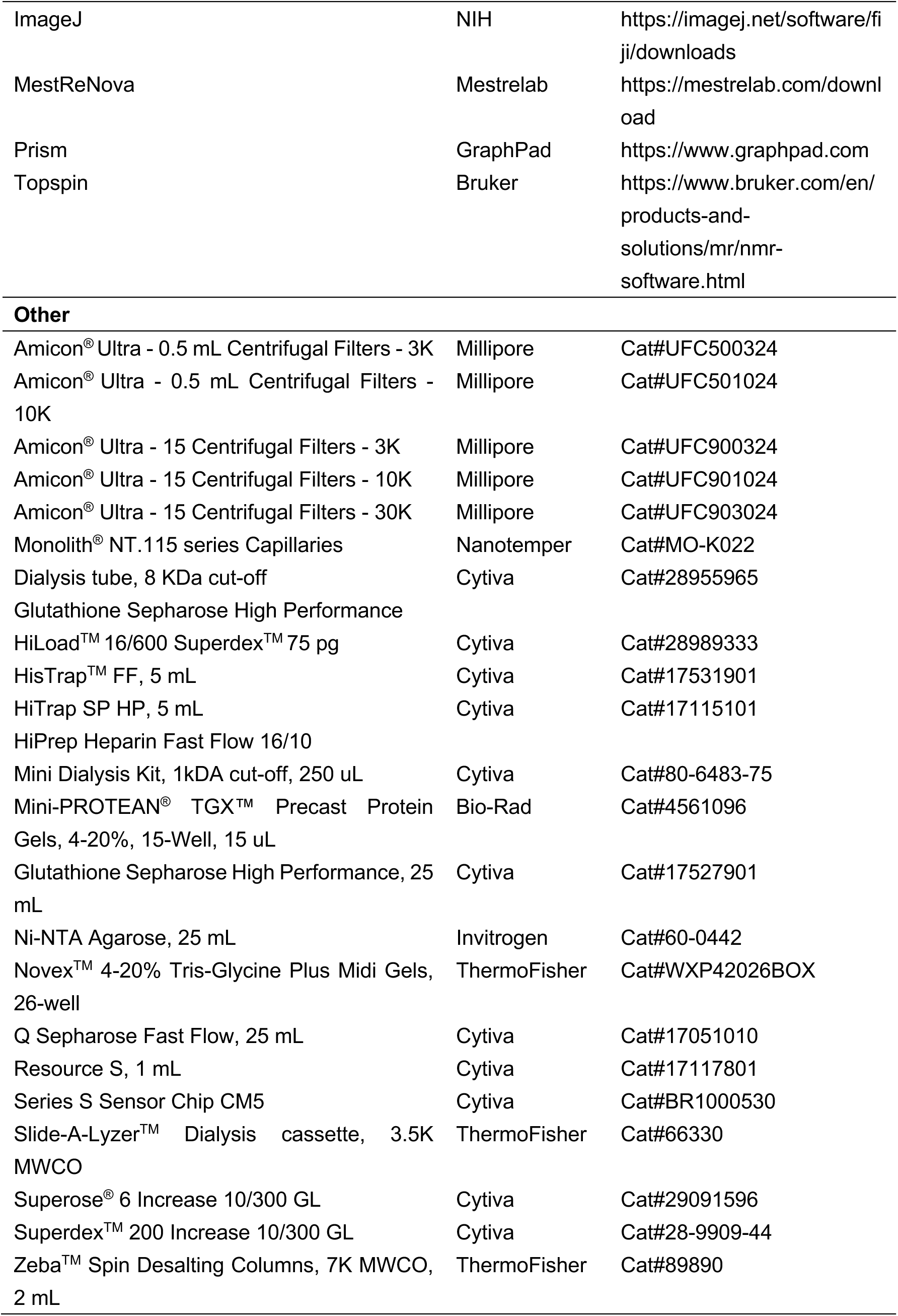
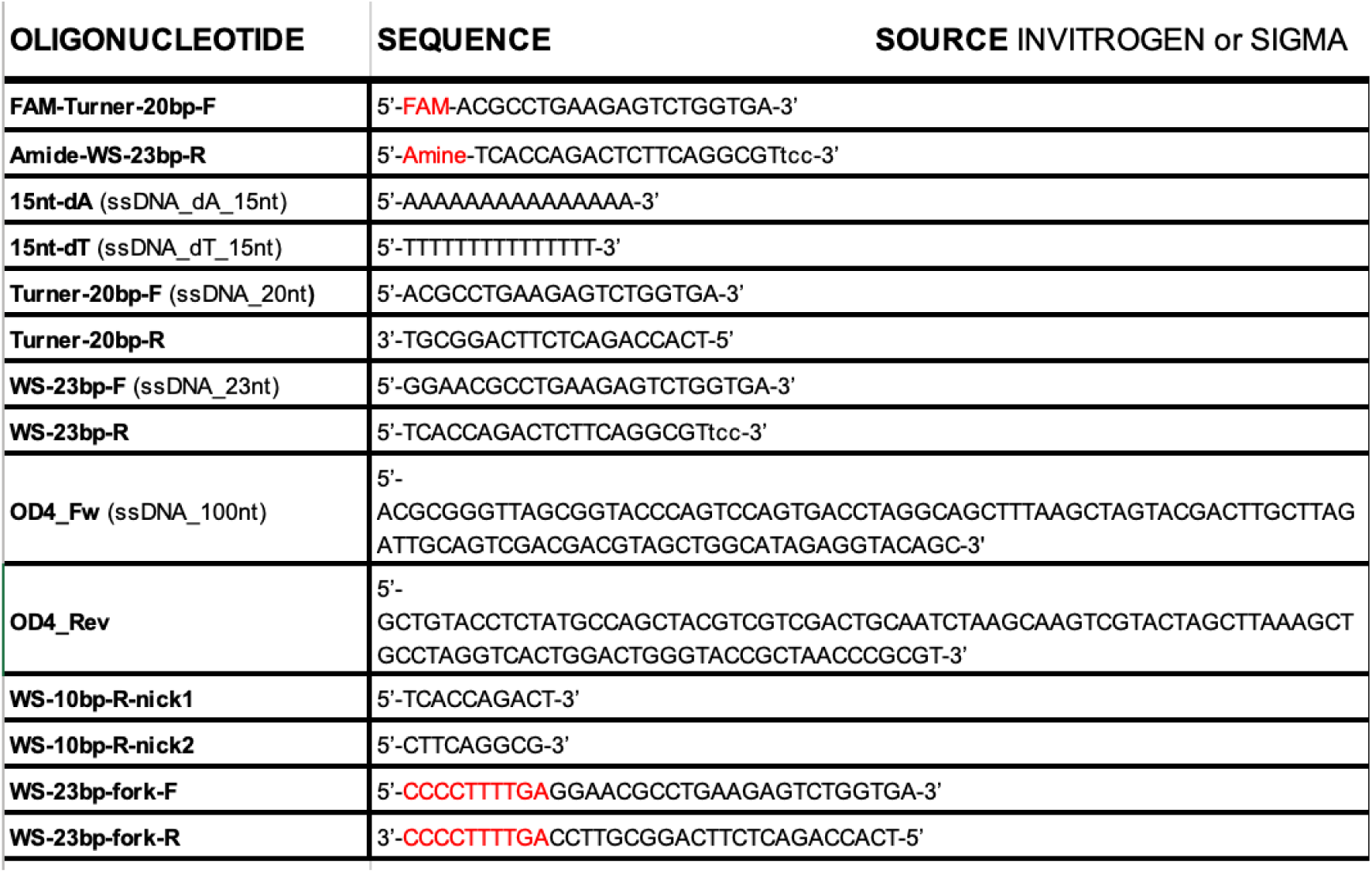
KEY RESOURCES TABLE.

### EXPERIMENTAL MODEL AND SUBJECT DETAILS

#### Cell culture

U2OS and Hek293 cells were grown in Dulbecco’s Modified Eagle’s medium (DMEM, Sigma-Aldrich) supplemented with 10% fetal bovine serum (Sigma-Aldrich) and 100 I.U./mL penicillin - 0.1 mg/mL streptomycin (Sigma-Aldrich) at 37°C in a humidified incubator with 5% CO_2_. Cells were tested monthly for mycoplasma contamination. SPRTN was depleted from U2OS with siRNA (si3’UTR: 5’-GUCAGGAAGUUCUGGUUAA-3’, Mycrosynth) for 3 days before processing. Formaldehyde treatment in Hek293 cells was performed in complete DMEM medium for 30 minutes at 37°C.

### METHOD DETAILS

#### DPC isolation and detection

DPCs were isolated using a modified rapid approach.^35,38^ To detect H1.0 crosslinks, the equivalent of 15 µg of the isolated DNA was digested with benzonase (50 units, 1 hour at 37°C) in a total of 400 µl. The entire volume was loaded onto a slot blot manifold (Biorad) and applied to a nitrocellulose membrane. The membrane was processed for western blot against H1.0 (Proteintech, 17510-1-AP). Alternatively, 80 µg of DNA were digested with benzonase (50 units, 1 hour at 37°C) in a total of 1 mL; proteins were precipitated with Trichloroacetic Acid (TCA) and resolved by SDS-. After transfer on nitrocellulose and western blot, H1.0 DPCs were detected with the H1.0 antibody. For slot blot detection of dsDNA, 100-200 ng of DNA were incubated with 50 µg/ml proteinase K to digest the crosslinked proteins, diluted in Tris/Borate/EDTA (TBE) buffer and applied to a nylon membrane (GE Healthcare). After the western blot, DNA was detected with an anti-dsDNA antibody (Abcam, ab27156).

### Purification of recombinant proteins

#### Ubiquitin purification

To express and purify monoUb, the *E.coli* BL21(DE3) strain carrying the pGEX-6P1-hUb plasmid was grown in LB medium at 37 °C with constant shaking until OD_600_ reached ∼0.9. It was induced by IPTG at a final concentration of 0.2 mM at 20 °C overnight. The harvested cell pellet was resuspended in ∼50 mL GST Buffer (50 mM Tris-HCl, pH7.4, 300 mM NaCl, 1 mM DTT) containing 0.2 mM PMSF and one tablet EDTA-free protease inhibitor cocktail (ROCHE) and then processed by sonication. The lysate was centrifuged at 20K for 30 min. The above supernatant was loaded onto the pre-equilibrated self-packed column with 10 mL Glutathione Sepharose High Performance (Cytiva). Then, the column was washed with GST Buffer until the UV became steady. The protein was eluted with GST Buffer containing 20 mM reducing GSH. The elution was added with HRV 3C protease for overnight cleavage at 4 °C using a 3 mL 3500 MWCO Slide-A-Lyzer Dialysis cassette (ThermoFisher) against 500 mL GST Buffer. The above O/N cleavage sample was loaded onto the GSH column to separate free GST from the tag-free monoUb. The flowthrough was collected and concentrated for SEC by HiLoad^TM^ 16/600 Superdex^TM^ 75 pg with the SEC Buffer (50 mM HEPES, pH 7.4, 150 mM NaCl, 0.5 mM TCEP-HCl). The tag-free NEDD8 was expressed by pGEX2TK-NEDD8 following the same procedure above.

To express and purify His-tagged monoUb/M1-Ub_2_/M1-Ub_4_/M1-Ub_6_ (including variants), *E.coli* BL21(DE3) strain carrying the corresponding plasmid was grown in LB medium at 37 °C with constant shaking until OD_600_ reached to ∼0.9. It was then induced by IPTG at a final concentration of 0.2 mM at 20 °C overnight. The harvested cell pellet was resuspended in Equilibration Buffer (50 mM HEPES, pH 7.4, 300 mM NaCl, 1 mM TCEP-HCl) with additionally 0.2 mM PMSF and 30 mM imidazole and processed by sonication. The lysate was centrifuged at 20 K rpm for 30 min. The supernatant was loaded onto the pre-equilibrated HisTrap FF column (Cytiva), and then the mixture was washed with Equilibration Buffer containing 30 mM imidazole until UV became steady. Then, the protein was eluted with an Equilibration Buffer containing 300 mM imidazole. Elution was concentrated for the final SEC purification via the HiLoad^TM^ 16/600 Superdex^TM^ 75 pg column with the SEC Buffer (50 mM HEPES, pH 7.4, 150 mM NaCl, 0.5 mM TCEP-HCl).

To express N^15^-labelled His-tagged monoUb, *E.coli* BL21(DE3) strain carrying the pNIC28-Ub plasmid was grown in N^15^-M9 medium (containing 6 g/L Na_2_HPO_4_, 3 g/L KH_2_PO_4_, 0.5 g/L NaCl, 2g/L D-glucose, 0.7 g/L ^15^NH_4_Cl, 1X gibco® MEM vitamin solution, 0.1 M FeSO_4_, 1M CaCl_2_, 1M MgSO_4_, pH 7.4) at 37 °C with constant shaking until OD_600_ reached to ∼1.0. It was then induced by IPTG at a final concentration of 0.2 mM at 20 °C overnight. The purification procedure is the same as the unlabelled Ubs from above except that the SEC purification was performed with NMR Buffer (22 mM phosphate, pH 7.0, 55 mM NaCl, 1 mM DTT). N^15^-labelled M1-Ub_2_ was expressed and purified in a similar way with the addition of HRV 3C protease for overnight cleavage to remove the His tag.

#### SPRTN purification

SPRTN FL was expressed and purified following the previously published protocol.^3^ To express GST-SPRTN-ΔC, *E.coli* Rosetta (DE3) strain carrying the pGEX-4T1-SPRTN-ΔC (Lys241AsnfsX8) plasmid was grown in LB medium (containing 100 µM ZnCl_2_) at 37 °C with constant shaking until OD_600_ reached to ∼0.8. It was induced by IPTG at a final concentration of 0.2 mM at 18 °C overnight. The harvested cell pellet was resuspended in ∼50 mL GST High Salt Buffer (100 mM HEPES, pH7.4, 500 mM NaCl, 10% Glycerol, 1 mM DTT) containing 0.2 mM PMSF, 0.1% DDM, 45 U/mL benzonase and one tablet EDTA-free protease inhibitor cocktail (Roche) and then processed by sonication. The lysate was centrifuged at 20K for 30 min. The purification procedure is similar to that of GST-Ub, except that GST High Salt Buffer was used to equilibrate the column, and protein was eluted by GST High Salt Buffer, which contained 20 mM of reduced GSH. The GST tag was not removed. The following SEC purification was performed by HiLoad^TM^ 16/600 Superdex^TM^ 75 pg with the GST SEC Buffer (50 mM HEPES, pH 7.4, 500 mM NaCl, 10% Glycerol, 1 mM DTT). GST-MIU was purified similarly, except that the Buffer used for SEC purification is 50 mM HEPES, pH 7.4, 150 mM NaCl, and 0.5 mM TCEP-HCl. SPRTN ZBD-BR was purified in a similar way in addition to HRV 3C protease to remove the GST tag. Tag-free SPRTN ZBD-BR was purified by SEC with SPRTN Buffer (50 mM HEPES, pH 7.4, 500 mM NaCl, 1 mM MgCl_2_, 10% Glycerol, 1 mM TCEP-HCl).

To express the SPRTN core (including the L38A, L99A and E112Q variants), *E.coli* BL21(DE3) strain carrying pNIC28-SPRTN (26-240) plasmid was grown in LB medium (containing 0.1 mM ZnCl_2_) at 37 °C with constant shaking until OD_600_ reached to ∼1.0. It was then induced by IPTG at a final concentration of 0.5 mM at 18 °C overnight. The harvested cell pellet was resuspended in SPRTN Buffer containing 45 U/mL benzonase, 0.04 mg/mL Pefabloc, one tablet EDTA-free protease inhibitor cocktail (Roche), and processed by sonication. The lysate was centrifuged at 20 K rpm for 30 min. The supernatant was loaded onto the pre-equilibrated self-packed column with 20 mL Ni-NTA resin (Invitrogen) and washed with SPRTN Buffer containing 30 mM imidazole until UV became steady. Then the protein was eluted with SPRTN Buffer containing 300 mM imidazole. Elution was concentrated and added with TEV protease to remove the tag. At the same time, it was dialysed overnight against 0.5 L SPRTN Buffer using an 8 kDa cut-off dialysis tube (Cytiva). The dialyzed sample from TEV cleavage was loaded onto the pre-equilibrated 5 mL HisTrap^TM^ FF crude (Cytiva) by SPRTN Buffer. The flowthrough was collected and further concentrated for the final purification by SEC. SEC was performed by loading the protein onto the HiLoad^TM^ 16/600 Superdex^TM^ 75 pg column with the SPRTN Buffer.

#### H1 purification

All the H1 proteins were expressed by the pCold cold shock system. *E.coli* Rosetta (DE3) strain carrying the corresponding plasmid was grown in LB medium at 37 °C with constant shaking until OD_600_ reached ∼0.8. The culture was cold-shocked at 15 °C for 30 min without shaking. It was induced by IPTG at a final concentration of 0.5 mM at 15 °C for ∼ 24 h. The harvested cell pellet was resuspended in His-Eq Buffer (50 mM Tris-HCl, pH7.4, 300 mM NaCl, 10% Glycerol, 0.5 mM TCEP-HCl) containing 0.2 mM PMSF, 30 mM imidazole, 1 tablet EDTA-free protease inhibitor cocktail (Roche) and then processed by sonication. The lysate was centrifuged at 20K for 30 min. The supernatant was loaded onto the pre-equilibrated HisTrap FF column (Cytiva) (Cytiva), and then was washed with His-Eq Buffer containing 30 mM imidazole until UV became steady. H1 was eluted with His-Eq Buffer containing 300 mM imidazole and further dialyzed into His-IEX Buffer (50 mM HEPES, pH 7.4, 150 mM NaCl, 10% Glycerol, 0.5 mM DTT). It was loaded onto the pre-equilibrated HiTrap SP column (Cytiva). H1 was eluted using a linear 10 CV gradient of 100 mM to 1.5M NaCl. The pure H1 fractions were pooled and dialyzed into His-Eq Buffer to reduce salt strength. Ub-H1, M1-Ub_4_-H1 and 2xISG15-H1 were purified in the same way with additional SEC purification by Superose^®^ 6 Increase 10/300 GL (Cytiva) with His-Eq Buffer.

#### PCNA purification

To express and purify PCNA, the *E.coli* BL21(DE3) strain carrying the pET16b-PCNA plasmid was grown in TB medium at 37 °C with constant shaking until OD_600_ reached ∼0.7. It was then induced by IPTG at a final concentration of 0.5 mM at 16 °C overnight. The harvested cell pellet was resuspended in 50 mM Tris-HCl pH 8.0, 100 mM NaCl, 10% glycerol, 1 mM EDTA and 1 tablet EDTA-free protease inhibitor cocktail (Roche) and lysed by sonication. The lysate was centrifuged at 20K for 30 min. The supernatant was applied to Q Sepharose Fast Flow column (GE Healthcare) pre-equilibrated in Buffer A (20 mM Tris-HCl pH 8.0, 10% glycerol). The column was then washed with Buffer A until the UV became steady. Protein was eluted using a linear 8 CV gradient of 100 mM to 1.5M NaCl. Fractions containing PCNA were dialyzed into a Buffer B (20 mM Tris-HCl pH 8.0, 100 mM NaCl, 10% glycerol) and loaded onto the pre-equilibrated HiPrep Heparin Fast Flow 16/10 (GE Healthcare). The column was washed with 2 CV Buffer B, followed by a 5 CV elution gradient from 0% to 100% of Buffer B containing 1.5 M NaCl. Fractions containing PCNA proteins were collected and concentrated for SEC by Superdex 200 Increase 10/300 GL (Cytiva) with GF buffer (20mM Tris pH 8.0, 100 mM NaCl, 2.5 % Glycerol and 1 mM TCEP).

#### Cy5 labelling

To label naked H1 (Cys was introduced to either N- or C-terminal) with Cy5, ∼ 0.5 mg His-H1 was added with 10 uL Cy5 (one pack of Cy5 was dissolved in 50 uL DMSO, cytiva) in a 200 uL total reaction with Labelling Buffer (50 mM HEPES, pH 7.4, 300 mM NaCl, 0.5 mM TCEP-HCl). React at R.T. in the dark for ∼2.5h. The labelled protein was separated from the free dye by SEC using the Superose^®^ 6 Increase 10/300 GL (Cytiva) with the Labelling Buffer. Peak fractions were collected and concentrated/buffer-exchanged with new SEC Buffer by a 10 K mini-concentrator (Millipore) to remove any residue-free dye. The reaction for labelling of the Ub-fused H1 proteins was prepared the same as above. React in the dark at 4 °C for ∼2h. Due to the low stability of the Ub-fused H1 proteins, after incubation, the reaction was directly concentrated/buffer-exchanged with a new SEC Buffer several times by a 10 K mini-concentrator (Millipore) to remove any free dye.

To label SPRTN core (E112Q) with Cy5 for the MST measurements, 100 uL SPRTN core (691.5 µM) was mixed with 10 uL Cy5 (Cytiva, one vial was dissolved in 50 uL DMSO) in 90 uL SEC Buffer (50 mM HEPES, pH 7.4, 150 mM NaCl, 0.5 mM TCEP-HCl). Incubate overnight in the dark at 4 °C. The labelled protein was separated from the free dye by SEC using the Superose^®^ 6 Increase 10/300 GL (Cytiva) with the SEC Buffer. Peak fractions were collected and concentrated/buffer-exchanged with new SEC Buffer by a 10K mini-concentrator (Millipore) to remove any residue-free dye.

#### Making dsDNAs

The ssDNAs (sequence provided in Key Resources Table) were diluted in DEPC-treated water to a final concentration of 100 µM. To make dsDNA, an equal volume of ssDNA was mixed and incubated at 95 °C for 10 min on a heating block. The mixture was then left on the heating block switched off to allow cooling down with gradient temperature drop naturally until equilibrium to room temperature. dsDNA_dAT_15nt was made with 15nt-dA and 15nt-dT; dsDNA_20nt was made with Turner-20bp-F and Turner-20bp-R; dsDNA_23nt was made with WS-23bp-F and WS-23bp-R; dsDNA_100nt was made with OD4_Fw and OD4_Rev; dsDNA_20/23nt was made with Turner-20bp-F and WS-23bp-R; dsDNA_20nt_nicked was made with Turner-20bp-F, WS-10bp-R-nick1 and WS-10bp-R-nick2; dsDNA_23/10bp_fork was made with WS-23bp-fork-F and WS-23bp-fork-R.

#### Making model H1-DPCs

The FAM-labelled dsDNA_20/23nt (FAM-Turner-20bp-F with Amide-WS-23bp-R, sequence provided in Key Resources Table) was produced by annealing following the above protocol and then purified using a Zeba Spin desalting column (7 kDa, 3 mL, Thermo Scientific, 89889) pre-equilibrated with Crosslink Buffer (100 mM NaH_2_PO_4_, 150 mM NaCl, 5 mM EDTA, pH7.3) to remove any unbound ssDNA. The flow-through was collected and further incubated with SMCC at a final concentration of 5 mM at R.T. in the dark for 2 h. Any excess SMCC was removed by a new Zeba Spin desalting column. The flowthrough containing dsDNA-SMCC was collected for conjugating with proteins (H1, Ub_4_-H1). 5-fold excess of dsDNA-SMCC was mixed with the protein and incubated overnight at 4 °C in the dark. The mixture was applied to IEX by a 1 mL Resource S column (Cytiva) pre-equilibrated with IEX Buffer A (50mM HEPES pH7.4, 150mM NaCl, 10% Glycerol, 0.5mM DTT). The protein conjugated to dsDNA was eluted by IEX buffer B (50mM HEPES pH7.4, 1M NaCl, 10% Glycerol, 0.5mM DTT) and concentrated by a 3K mini-concentrator (Millipore) as the final DPC product.

#### Making Ub-PCNA

Monoubiquitination of PCNA *in vitro* was adopted from a previously published protocol,^62^, with below optimization. Ube1 (1 µM), Ubc5Hc-S22R (10 µM), His-monoUb (30 µM), PCNA (2 µM) and ATP (10 mM) were prepared in MMT Working Buffer (50 mM MMT, pH 9, 25 mM NaCl, 3 mM MgCl_2_, 0.5 mM TCEP-HCl) and incubated overnight at 30 °C. The sample was centrifuged where the supernatant was loaded onto the Superose^®^ 6 Increase 10/300 GL (Cytiva) with SEC Buffer (50 mM HEPES, pH 7.4, 150 mM NaCl, 0.5 mM TCEP-HCl) to separate Ub-PCNA from other ubiquitination components.

#### SPRTN cleavage assay

For the SPRTN cleavage towards free H1 substrates, recombinant SPRTN (or variants) (2 µM) were incubated with H1 (1 µM) in the absence or presence of dsDNA_20/23nt (2.7 µM) in combination with Ubs or Ubs (2 µM, unless specified otherwise) with the indicated time at 30°C. For the SPRTN cleavage towards H1-DPC substrates, Recombinant SPRTN (or SPRTN core) (2 µM) were incubated with H1-DPC substrates (∼0.1 µM) in combination with M1-Ub_4_ (2 µM) with the indicated time at 30°C. Details can be found in each corresponding figure legend.

#### Microscale Thermophoresis (MST)

A serial dilution of monoUb (from 4754.8 µM till 0.15 µM) and 1.2 µM Cy5-labelled SPRTN core were prepared in a total volume of 20.5 uL with MST Buffer (50 mM HEPES, 150 mM NaCl, 0.5 mM TCEP-HCl). Each sample was loaded to the capillary (Nanotemper) for the measurement with 20% MST Power on a microscale thermophoresis instrument (Monolith NT.115, Nanotemper). The data were analyzed and plotted by the NT Analysis Software (1.5.41, Nanotemper). All the analyses were done using the “Thermophoresis + Temperature Jump” method. The fluorescent value is fixed to 1.2 µM for the fitting.

#### Isothermal Titration Calorimetry (ITC)

Protein samples were dialysed against ITC buffer (50 mM HEPES pH 7.4, 50 mM NaCl, 0.5 mM TCEP-HCl) overnight at 4°C using the Mini Dialysis Kit (1 KDa cut-off, Cytiva) before measurements. All the ITC measurements were conducted in a MicroCal PEAQ-ITC calorimeter (Malvern) with a standard 13-injection titration program (1st injection: 0.4 μl/injection, duration: 0.8s, spacing time: 150 s; the rest of the injections: 3 μl/injection, duration: 6s, spacing time: 150 s) at 25°C, with a reference power at 10 μCal/s. Stir speed is 750 rpm. Initial delay is 60s. The data is processed and analyzed using the MicroCal PEAQ-ITC Analysis Software with a one-site binding model.

#### Surface Plasmon Resonance (SPR)

SPR measurements were performed on a Biacore^TM^ S200 (GE Healthcare). SPRTN PIP-UBZ was immobilized onto a Series S Sensor sensor chip CM5 (Cytiva) by an amine coupling kit (Cytiva) at pH 5.0 until the response on the surface reached ∼ 1000 RU. A serial dilution of the analytes (monoUb, PCNA and Ub-PCNA) was prepared on a 96-well plate. Analytes were injected onto the chip at a flow rate of 35 μL/min with 60 s association and 200 s dissociation. The signal from the reference channel was subtracted from the signal obtained from the sample channel. The data were fitted using the Biacore S200 Evaluation software (GE Healthcare) using a steady state affinity binding model with report points taken at 8 s after injection start.

#### NMR spectroscopy

^15^N-labelled proteins (His-monoUb, M1-Ub_2_, SPRTN core) were prepared at a concentration of 100 µM in NMR Buffer (22 mM phosphate, pH 7.0, 55 mM NaCl, 1 mM DTT) containing 5% D_2_O and 0.05% NaZ. NMR spectra were recorded on a 950 MHz spectrometer (Bruker Avance III HD console) equipped with a high-sensitivity 5 mm TCl cryoprobe at 25 °C.

For the titration between ^15^N-labelled His-monoUb (or ^15^N-labelled M1-Ub_2_) and SPRTN core, a series of titrations are made by the addition of unlabelled SPRTN core to the ^15^N-labelled His-monoUb (or ^15^N-labelled M1-Ub_2_). HSQC Spectra were recorded. Spectra were initially processed using TopSpin 3.6 and then analysed and plotted using MestReNova. The raw data document will provide all the quantitative analysis of the peak volume from the NMR titration (will be available on Mendeley). The volume of the peaks from monoUb or M1-Ub_2_ that are completely broadened upon the addition of SPRTN core or UBZ is defined as 0.

#### Size-exclusion chromatography coupled multiple-angle laser light scattering (SEC-MALS)

UBZ or UBZ* were prepared at 3.0 mg/mL and then 100 uL of each concentration was separately applied to the pe-equilibrated SEC column (Superdex® 75 10/300 Increase, Cytiva) using the buffer containing 50 mM HEPES pH 7.4, 150 mM NaCl, 0.5 mM TCEP-HCl. The protein eluted from SEC was monitored using a DAWN HELEOS-II 18-angle light scattering detector (Wyatt Technologies), a U9-M UV/Vis detector (Cytiva), and an Optilab T-rEX refractive index monitor (Wyatt Technologies). Data were analysed by using Astra v7 (Wyatt Technologies) with a refractive increment value of 0.185 mL/g.

#### Data analysis

Multiple sequence alignment was performed by CLUSTAL (1.2.4).

For the kinetic data from the SPRTN cleavage assay, the signal from each time point was normalized to the “0 min” time point within each condition. Kinetic data were fitted with simple linear regression or one-phase exponential decay - least squares fit (Prism). Significant analysis from SPRTN cleavage assay was performed using a paired t-test (Prism). *p <0.05; **p <0.005; ***p <0.0005. Details can be found in each corresponding figure legend.

